# The Kaposi’s sarcoma-associated herpesvirus viral genome packaging accessory factor ORF68 forms cytoplasmic puncta dependent on the viral tyrosine kinase

**DOI:** 10.64898/2026.02.23.707506

**Authors:** Sara Gelles-Watnick, Dawei Liu, Mayte A. Cerezo-Matias, Jessica Liu, Giancarlo Ospina, Alicia Schäfer, Lilian Kabeche, Ileana M. Cristea, Allison L. Didychuk

## Abstract

Assembly of Kaposi’s sarcoma-associated herpesvirus (KSHV) virions is dependent on ORF68, a viral genome packaging accessory factor of unknown function. We used confocal fluorescence imaging to monitor ORF68 localization throughout the KSHV lytic cycle. ORF68 localizes to nuclear viral replication compartments, the site of viral genome packaging. Unexpectedly, ORF68 also localized to cytoplasmic puncta. Using proximity labeling mass spectrometry during infection, we identified ORF68 interaction partners, including the viral tyrosine kinase ORF21. We show that ORF21 colocalizes with ORF68 in cytoplasmic puncta and draws ORF68 to the cytoplasm during infection. This interaction is mediated by the disordered N-terminal region of ORF21. We propose that ORF68 possesses a novel secondary function independent from virion assembly.

**IMPORTANCE:** Kaposi’s sarcoma-associated herpesvirus (KSHV) is the underlying cause of multiple human malignancies including Kaposi’s sarcoma, primary effusion lymphoma, and multicentric Castleman’s disease. Encapsidation of the viral genome is necessary to produce new infectious virus. Viral genome packaging can be targeted with small molecules in human cytomegalovirus, a related herpesvirus, but we lack a sufficiently detailed mechanistic understanding to develop additional therapeutics. The molecular role of the essential packaging accessory factor, encoded by ORF68 in KSHV, has remained unclear. We studied the interactions and localization of ORF68 during KSHV infection and find that, in addition to its nuclear role in packaging, ORF68 forms cytoplasmic puncta through its interaction with the viral tyrosine kinase ORF21. This study demonstrates how conserved, essential herpesvirus proteins can play multiple roles in distinct subcellular compartments during viral infection, with secondary functions that may be herpesvirus species-specific.

## INTRODUCTION

Kaposi’s sarcoma-associated herpesvirus (KSHV) is an oncogenic human virus, responsible for cancers and malignancies including Kaposi’s sarcoma (KS), primary effusion lymphoma, and multicentric Castleman’s disease^1– 3^. Patients infected with HIV-1 and other immunosuppressive disorders are at high risk for KSHV infection with progression to KS^3,4^. Like all herpesviruses, KSHV is an enveloped, double-stranded DNA virus that cycles between latency and a lytic phase wherein daughter progeny virions are produced. There is no cure or prophylactic vaccine for KSHV and the therapeutic regimens used to treat other human herpesvirus infections have limited efficacy against KSHV^5^.

During lytic infection, the virally encoded DNA replication machinery synthesizes new viral genomes in the nucleus of infected cells^6–9^. Replication occurs in part through rolling circle replication, resulting in head-to-tail concatemers of the viral genome separated by terminal repeat sequences^10–17^. Viral genome replication results in a condensate within the nucleus called a viral “replication compartment” (vRC)^6,18–20^. Capsid components are expressed from newly replicated genomes, whereupon they are imported into the nucleus and assembled into icosahedrons where one unique vertex is occupied by the portal protein^21^. Viral genomes are encapsidated by a powerful viral packaging motor that cleaves within the terminal repeats to produce a capsid containing a full-length viral genome^22–24^. Herpesvirus genome packaging is thought to occur in vRCs^25–30^. Subsequently, the genome-packed capsid egresses from the nucleus, acquires a tegument layer, and undergoes secondary envelopment prior to release of progeny virions.

Six viral proteins conserved across the *Herpesviridae* are required for viral genome cleavage and packaging^31^. In KSHV, these proteins are encoded by ORF7, ORF29, ORF32, ORF43, ORF67A, and ORF68. Loss of these factors results in the accumulation of unpackaged, empty “B capsids” in nuclei of infected cells^24,29,31–34^. The role of five of these components is clearly defined. The viral packaging motor – called the terminase – consists of ORF7, ORF29, and ORF67A^24,32,33,35^. The portal protein, ORF43, is required for viral DNA entry and exit from the capsid^34,36^, while ORF32 is a component of the capsid-associated tegument complex^37,38^.

Homologs of the sixth component, ORF68, are required in all three herpesvirus subfamilies, and loss of function or expression prevents infectious virion production^29,39–42^. However, the mechanistic contribution of ORF68 and its homologs to viral genome packaging remains unclear. High-resolution structures of KSHV capsids indicate that ORF68 is not a constitutive component of the capsid^38,43,44^. Structural and biochemical characterization of ORF68 show that it assembles into a homopentameric ring with a positively charged central channel that is critical for viral genome packaging^40^. Little is known about ORF68 protein-protein interactions or subcellular localization during the KSHV lytic phase. Characterization in the context of viral infection is needed to further elucidate the precise function of ORF68.

In this study, we used confocal microscopy to visualize ORF68 during KSHV lytic replication and found that ORF68 forms cytoplasmic puncta in addition to its canonical localization to the nucleus for viral genome packaging. These cytoplasmic puncta lack features consistent with known cellular condensates or organelles and do not colocalize with other components of the viral packaging machinery. Using biotinylation-based proximity labeling, we identified binding partners of ORF68, including the viral tyrosine kinase ORF21. We demonstrated that ORF21 colocalizes with ORF68 cytoplasmic puncta and that ORF21 expression is required for ORF68 cytoplasmic localization during viral infection. Finally, we show that the intrinsically disordered N-terminal region of ORF21 is necessary to drive ORF68 puncta formation independent of viral infection. Thus, we have shown that the ORF21 – ORF68 interaction regulates the partitioning of ORF68 to cytoplasmic puncta during the lytic cycle, potentially for a function separate from nuclear viral genome packaging.

## RESULTS

### ORF68 tolerates addition of an N-terminal tag during viral replication

We engineered a fluorescent monomeric NeonGreen (mNG) tag at the endogenous ORF68 locus in the KSHV genome to track the subcellular localization of ORF68 throughout viral replication. We used Red recombineering to modify KSHV BAC16 such that the vector backbone eGFP cassette was replaced with monomeric infrared fluorescent protein (mIFP) and mNG was fused to the N-terminus of ORF68 (**Fig. 1A, Fig. S1A**)^45^. As a control, we generated an additional virus where a T2A ribosome skipping sequence was inserted between mNG and ORF68 (mNG-T2A-ORF68) so that mNG and ORF68 were expressed separately^46^. We used the engineered BACs to establish latently infected iSLK cell lines^45,47,48^. We chemically reactivated the iSLK cell lines with doxycycline and sodium butyrate to monitor viral progression throughout the lytic cycle. Fusion of mNG to ORF68 did not affect ORF68 expression or expression of representative early (ORF6) or late (K8.1) viral genes (**Fig. 1B**). Using qPCR, we monitored release of encapsidated virions and found that addition of mNG or mNG-T2A to the ORF68 locus did not disrupt the production of progeny virion (**Fig. 1C**). Thus, addition of a large N-terminal fusion does not disrupt the essential function of ORF68 during KSHV replication.

**FIGURE 1.**
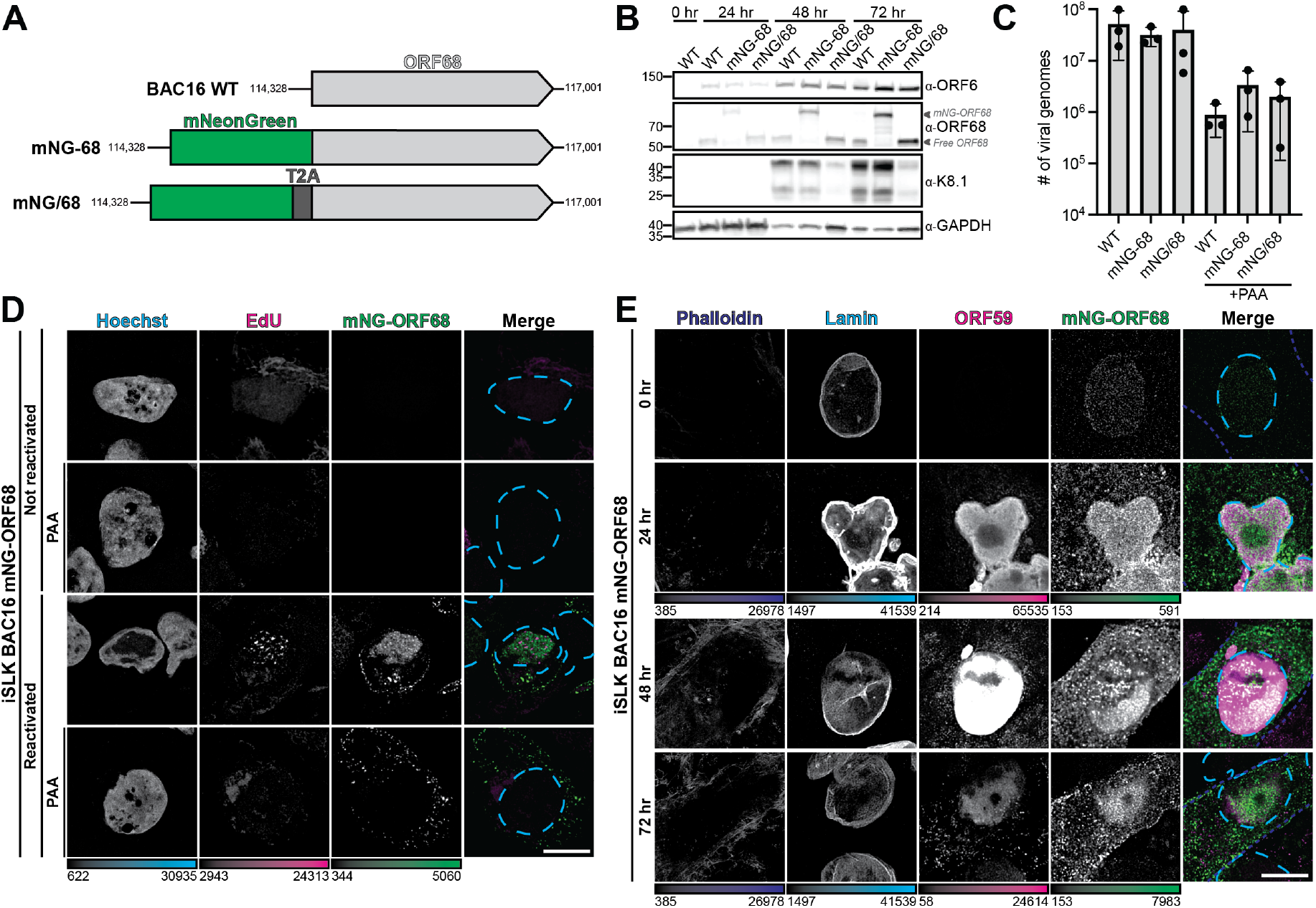
ORF68 localizes to viral replication compartments in the nucleus and non-spherical puncta in the cytoplasm. **A**. Schematic of the ORF68 locus in the BAC16 WT genome and insertion of Strep-tag (Strep-68), mNeonGreen (mNG-68) and mNeonGreen-T2A (mNG/68). **B**. Western blot of whole cell lysate (18 μg) from reactivated iSLK cells (0, 24, 48, or 72 hours post-reactivation) infected with BAC16 WT, mNG-ORF68, or mNG/68 virus, representative of three biological replicates. Representative early (ORF6) and late (K8.1) proteins are used to show progression through the virus lifecycle. GAPDH is the loading control. **C**. Extracellular virion production of reactivated iSLK cells infected with BAC16 WT, mNG/68, and mNG-68, with or without phosphonoacetic acid (PAA) treatment was quantified by qPCR from three biological replicates. Filled circles indicate the number of viral genome copies measured for each biological replicate, with the bar height representing the mean and error bars depicting the SD. None of the samples are significantly different from each other by pairwise RM one-way ANOVA (p>0.05). **D**. Z-slices of iSLK mNG-ORF68 cells at 72 hours post-reactivation with or without PAA treatment, representative of at least 10 cells from 2 biological replicates. Cells were incubated with EdU for 2 hours prior to fixation and click-conjugated to a AlexaFluor568 (magenta) and stained with Hoechst (cyan dashes). **E**. Z-slices of iSLK cells expressing mNG-ORF68 (green) at 0, 24, 48, and 72 hours post reactivation, representative of at least 10 cells from 2 biological replicates. Cells were stained for phalloidin (blue dashes), Lamin A/C (cyan dashes), and ORF59 (magenta). Scale bars are 10 μm; color bars represent the modified minimum and maximum pixel intensity values.

### ORF68 localizes to viral replication compartments in the nucleus, but forms puncta in the cytoplasm

We examined the subcellular localization of mNG-ORF68 throughout the lytic cycle using confocal microscopy. We stained host DNA with Hoechst and used EdU-Click chemistry to label nascently replicated viral genomes with a fluorescent probe. During viral lytic replication, changes to the cell cycle eliminate cellular DNA replication, while robust viral DNA replication forms EdU foci within vRCs^49,50^. Late in infection, mNG-ORF68 localizes to the nucleus, consistent with prior immunofluorescence (IF) microscopy examining ORF68 localization in iSLKs using an anti-ORF68 polyclonal antibody (**Fig. 1D**)^29^. Nuclear ORF68 does not colocalize with nucleolin, a marker for nucleoli (**Fig. S1B**). ORF68 localizes to Hoechst-negative nuclear regions containing EdU-labeled regions, indicative of vRCs. Treatment with an inhibitor of viral genome replication, phosphonoacetic acid (PAA), eliminated nuclear EdU signal (**Fig. 1D**).

Unexpectedly, we also observed formation of ORF68 puncta in the cytoplasm, including upon treatment with PAA (**Fig. 1D**). We monitored ORF68 localization throughout the lytic cycle (0, 24, 48, and 72 hours post reactivation), staining with phalloidin to demarcate the cytoplasm, lamin A/C to mark the nuclear boundary, and ORF59 as a protein marker for vRCs^6^. Consistent with prior IF^29^, ORF68 colocalizes with ORF59 in the nucleus throughout lytic infection (**Fig. 1E**). Cytoplasmic mNG-ORF68 puncta are visible throughout the lytic cycle, observable as early as 24 hours post-reactivation. To ensure that puncta formation was not an artifact caused by fusion of ORF68 to mNG, we visualized the same markers in iSLK cells infected with the mNG-T2A-ORF68 virus. Free mNG protein is expressed throughout the lytic cycle and is diffuse in the nucleus and cytoplasm (**Fig. S1C**). To ensure that ORF68 localization is not altered by fusion to the ∼27 kDa mNG tag, we generated an iSLK-BAC16 line with a Twin-Strep tag fused to the N-terminus of ORF68 (**Fig. S1A**). Upon reactivation into the lytic cycle, Strep-ORF68 is expressed to similar levels as untagged ORF68 (**Fig. S1D**). We performed immunofluorescence with an anti-Strep tag antibody on lytic cells infected with Strep-ORF68 or WT virus as a control (**Fig. S1E,F**). Similarly to mNG-ORF68, Strep-ORF68 colocalizes with ORF59 in the nucleus and forms puncta in the cytoplasm (**Fig. S1F**). Thus, we find that ORF68 is enriched in vRCs during the lytic cycle, consistent with its role in viral genome packaging, but that it also forms discrete cytoplasmic puncta throughout the viral lifecycle.

### ORF68 cytoplasmic puncta do not colocalize with the viral terminase, capsids, or cellular markers of cytoplasmic organelles

We next wanted to understand if other KSHV viral genome packaging components form cytoplasmic puncta. We modified the mNG-ORF68 BAC by inserting a hemagglutinin (HA) tag at the endogenous loci of ORF67A (N-terminus), ORF29 (C-terminus), or ORF65 (C-terminus) and established latently infected iSLK lines (**Fig. S2A,B**). All three proteins could be detected by western blot, albeit with differences in expression (**Fig. S2C**). Infectious virion production was not impacted, suggesting that these essential proteins remain functional upon addition of small epitope tags (**Fig. S2D**). ORF68 colocalized with KSHV terminase subunits (ORF29 and ORF67A) in the nucleus (**Fig. 2A**). Neither of the terminase subunits showed substantial cytoplasmic localization and thus did not colocalize with cytoplasmic ORF68 puncta (**Fig. 2B**). The predominantly nuclear subcellular localization of terminase subunits is consistent with prior IF studies of ORF29 and ORF67A homologs in HSV-1 and HCMV^18,51^.

**FIGURE 2.**
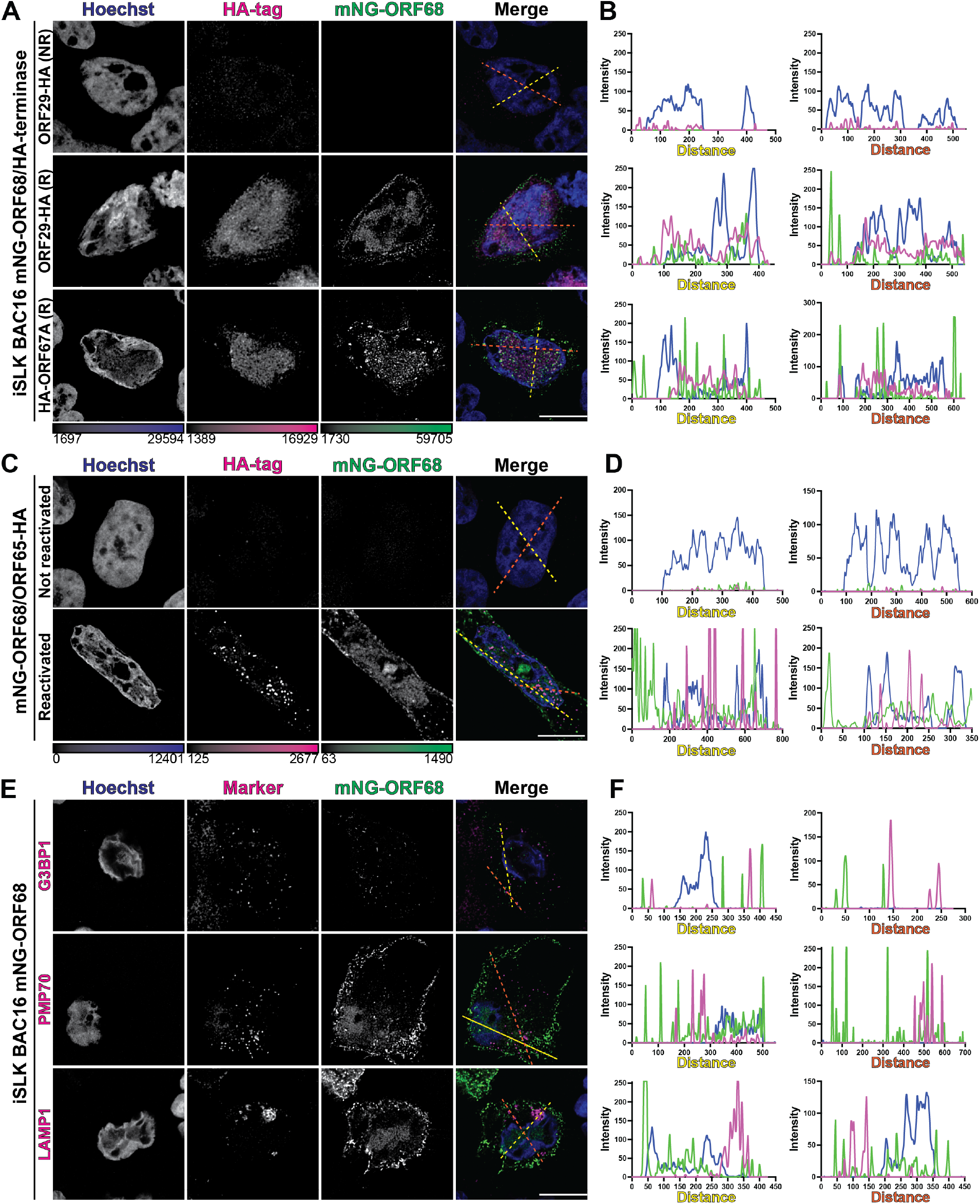
Cytoplasmic ORF68 puncta do not colocalize with viral packaging components nor with indicated markers of cytoplasmic organelles. **A**. Z-slices of lytic iSLKs expressing mNG-ORF68 (green) and either ORF29-HA or ORF67A-HA, fixed at 72 hours post-reactivation and stained for Hoechst (blue) and HA-tag (magenta). Representative of at least 10 cells from two biological replicates. **B**. Intensity profile plots showing intensity of Hoechst (blue), mNG-68 (green), or Ha-tag (magenta) signal at each point along a line (yellow or orange dashed line). **C**. Z-slices of lytic iSLKs expressing mNG-ORF68 (green) and ORF65-HA, fixed 72 hours post-reactivation and stained for Hoechst (blue) and HA-tag (magenta). Representative of at least 10 cells from two biological replicates. **D**. Intensity profile plots showing intensity of Hoechst (blue), mNG (green), or HA-tag (magenta) signal at each point along a line (yellow or orange dashed line). **E**. Z-slices of lytic iSLKs expressing mNG-ORF68 (green), fixed 72 hours-post reactivation and stained for Hoechst (blue) and markers (magenta) for stress granules (G3BP1), peroxisomes (PMP70), or lysosomes (LAMP1). Representative of at least 10 cells from two biological replicates. **E**. Intensity profile plots showing intensity of Hoechst (blue), mNG-68 (green), or organelle marker (magenta) signal at each point along a line (yellow or orange dashed line). Scale bars are 10 μm; color bars represent the modified minimum and maximum pixel values.

RF65, the small capsid protein, binds to major capsid protein hexons and, in KSHV, is required for capsid assembly and stability^52–54^. ORF65 was previously reported to be detectable in cytoplasmic puncta after reactivation in BCBL-1 cells^55^ and can be used to monitor incoming cytoplasmic capsids shortly after *de novo* infection in iSLK^36^. We observed that ORF65 localizes to vRCs, the site of capsid assembly and viral genome packaging, and forms cytoplasmic puncta consistent with capsids that have egressed from the nucleus (**Fig. 2C**). However, the ORF68 and ORF65 cytoplasmic puncta do not colocalize (**Fig. 2D**). This suggests that ORF68 is likely not responsible for chaperoning elements of the capsid into the nucleus for packaging, nor for chaperoning successfully packaged capsids out of the cell.

Induction or reduction of various cytoplasmic puncta plays an important role in KSHV infection. During infection, cGAS-DNA interactions induce condensate formation^56–59^ and G3BP1 condenses into stress granules in the cytoplasm^56,60–62^. These processes are related to cellular anti-viral defense and have not been implicated in viral packaging. However, many viral proteins serve dual functions, and we considered whether the ORF68 cytoplasmic puncta are tied to these previously established pathways by monitoring colocalization with cytoplasmic organelles or condensates. We reactivated mNG-ORF68 BAC16 iSLKs for 72 hours and performed IF with antibodies for G3BP1, PMP70, or LAMP1 proteins as markers for stress granules, p-bodies, peroxisomes, or lysosomes, respectively (**Fig. 2E**). ORF68 signal does not colocalize with any of these markers (**Fig. 2F**). Thus, while ORF68 colocalizes with essential packaging machinery in the nucleus, the cytoplasmic puncta may serve a secondary, unrelated function.

### Identification of ORF68-interacting proteins by proximity labeling mass spectrometry

Given that various host or viral markers failed to colocalize with ORF68, we pursued an unbiased approach to identify ORF68 interactions using proximity biotinylation labeling. We selected TurboID as a promiscuous biotin ligase because of its favorable labeling kinetics and high activity^63^, allowing us to identify transient protein-protein interactions that occur late in infection. We engineered BAC16 to insert TurboID (TID) at the N-terminus of the ORF68 locus (**Fig. 3A, S3A**). Given that viral genome packaging occurs in the nucleus, we also included a control where TID was expressed separated from ORF68 through inclusion of a T2A site and shuttled to the nucleus via an nuclear localization tag (NLS).

**FIGURE 3.**
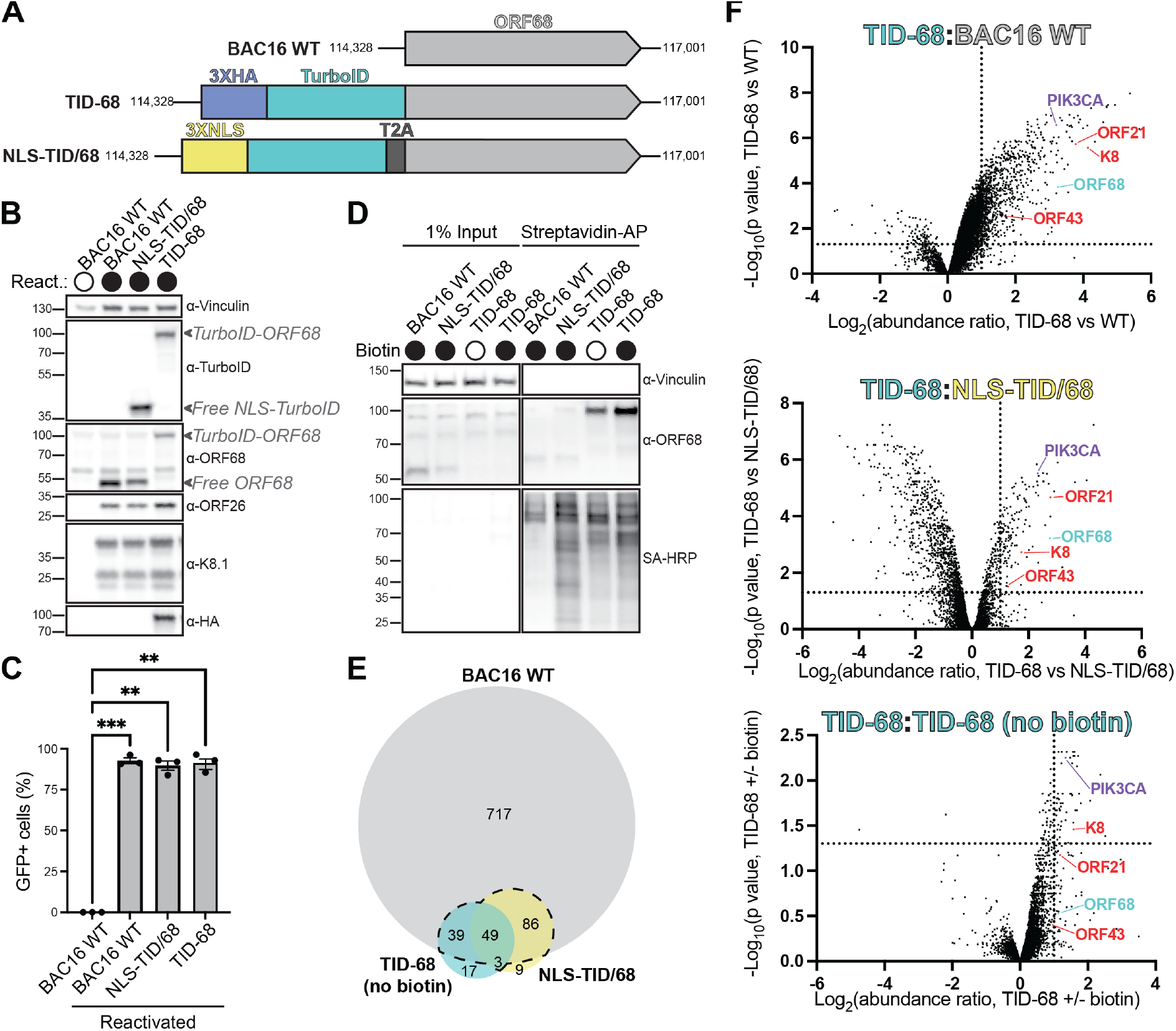
Proximity biotinylation identifies viral and host proteins that interact with ORF68. **A**. Schematic of BAC16 WT genome, showing position of ORF68 and the N-terminal insertions. Insertions include 3XNLS-TurboID-T2A (NLS-TID/68) and 3xHA-TurboID (TID-68). **B**. Western blot of whole cell lysate (25 μg) from reactivated WT, NLS-TID-T2A-ORF68, or TID-ORF68 iSLK cells 48 hours post-reactivation comparing protein levels of TurboID, HA-tagged and untagged ORF68, and representative late genes (K8.1, ORF26), representative of two biological replicates. Vinculin is the loading control. **C**. Supernatant transfer assay to assess infectious virion production from iSLK cells 72 hours post-reactivation. Data are from three biological replicates By repeated measure (RM) one-way ANOVA, there is a significant difference between the unreactivated BAC16 WT and all reactivated samples. ** indicates p-value of <0.005. *** indicates p-value of <0.001. **D**. Western blot of 4 mg streptavidin affinity purification of biotinylated proteins from iSLK cells 48 hours post-reactivation using Streptavidin-HRP (SA-HRP), representative of three biological replicates. Vinculin is the loading control. **E**. Venn diagram showing number of proteins enriched in the TID-68-expressing sample over WT, NLS-TID/68, and TID-68 (no biotin). **F**. Volcano plot comparing fold enrichment (x-axis) vs. p-value (y-axis) of proteins in the TID-68 sample compared to WT, NLS-TID/68, or TID-68 (no biotin) samples. The dotted lines represent enrichment ratio=2 (x-axis) and p-value=0.05 (y-axis). Enriched viral proteins are labeled in magenta, with select enriched host hits enriched in cyan.

We established iSLK cell lines latently infected with BAC16 WT, NLS-TID/68 (nuclear free TurboID control), or TID-ORF68 (**Fig. 3A, S3A**). Using an anti-TurboID antibody, we showed that free TID (from the NLS-TID/68 control) and TID-ORF68 are expressed at comparable levels (**Fig. 3B**). An anti-ORF68 polyclonal antibody also showed that ORF68 is expressed at comparable levels in all three cell lines and that T2A cleavage is complete (**Fig. 3B**). Late genes (K8.1, ORF26) were comparably expressed (**Fig. 3B**) and viral genome replication was similar between the three cell lines (**Fig. S3B**). Using a supernatant transfer assay, we found that infectious virion production was similar between WT, TID-68, and T2A viruses, indicating that addition of the TID tag at the ORF68 locus, like addition of the mNG tag, does not affect the production of progeny virions (**Fig. 3C**).

We performed proximity labeling 48 hours post-reactivation and used streptavidin affinity purification coupled with mass spectrometry to identify biotinylated proteins^63^. Proteins physically proximal (∼10 nm) to TurboID are covalently biotinylated during a 15-minute labeling period^64^. As controls, we included samples from cells infected with (1) WT virus (i.e., no TID) incubated with biotin to control for endogenously biotinylated proteins, (2) NLS-TID/68 virus (i.e., free nuclear TurboID) incubated with biotin to filter out non-specifically biotinylated nuclear proteins, and (3) TID-68 virus incubated without biotin to control for basal TurboID activity. As expected, the NLS-TID cell line exclusively biotinylated nuclear proteins, while biotinylated protein in TID-ORF68 cells localized to both the cytoplasm and nucleus (**Fig. S3C**). Biotinylation, followed by streptavidin affinity-purification and western blotting, indicated that expression of TID increased the total amount of biotinylated protein (**Fig. 3D**). Biotinylation of NLS-TID/68 and TID-ORF68 cell lines resulted in different banding patterns of biotinylated proteins, indicating that these proteins likely experience different interactions.

We next performed proximity labeling and used mass spectrometry to identify potential TID-protein interactions in each sample by processing three replicates of the four conditions (WT+biotin, TID-ORF68 no biotin, TID-ORF68+biotin, and NLS-TID/68+biotin) in parallel. Protein abundances across all samples were normalized to the abundance of the mitochondrial protein PCCA, which served as an internal constitutively biotinylated control, before differential abundance analysis^65^. We compared our TID-ORF68+biotin sample to the three controls and identified proteins with p-values <0.05 and fold-change >2. Many hits (891) were enriched over the WT control, while 147 and 108 hits were enriched over the NLS-TID/68 and TID-68 no biotin controls, respectively (**Fig. 3E, F**). We detected 49 proteins that were enriched over all three controls and a larger set of proteins (177) that were enriched in two of three controls, including ORF68 (**Supplementary Table S1**). Three additional viral proteins passed two of three cutoffs: K8, ORF43, and ORF21 (**Fig. 3F**). Gene ontology biological function analysis^66–68^ of the identified host proteins revealed that the majority of the hits function in cytoplasmic processes (**Fig. S3D**), consistent with our observations of ORF68 localization during infection. A consistent top cellular hit was the phosphoinositide 3-kinase 1-alpha (PI3K1α) catalytic subunit, p110α (PIK3CA) (**Fig. 3F**).

### ORF21 is required for cytoplasmic localization of ORF68 during infection

ORF21, a viral cytoplasmic protein that functions as a tyrosine kinase in KSHV infection^69–71^, was one of the most highly enriched proteins in the proximity labeling experiment. We sought to directly assess if ORF68 colocalizes with ORF21 in the cytoplasm of infected cells. We inserted an HA-tag at the C-terminus of ORF21 in the mNG-ORF68 BAC and generated a latently infected iSLK cell line, allowing us to simultaneously visualize both ORF68 and ORF21 (**Fig. S4A**). After 72 hours of lytic reactivation, we stained the cells for HA-tag, Lamin A/C, and phalloidin to demarcate ORF21, the nuclear envelope, or the cell membrane, respectively and visualized mNG-ORF68 (**Fig. 4A**). ORF21 colocalizes with ORF68 in cytoplasmic puncta at 72 hours post-reactivation (**Fig. 4B**). Colocalization of ORF68 and ORF21 in cytoplasmic puncta persisted upon treatment with PAA, suggesting that these proteins interact regardless of viral genome replication and capsid formation.

**FIGURE 4.**
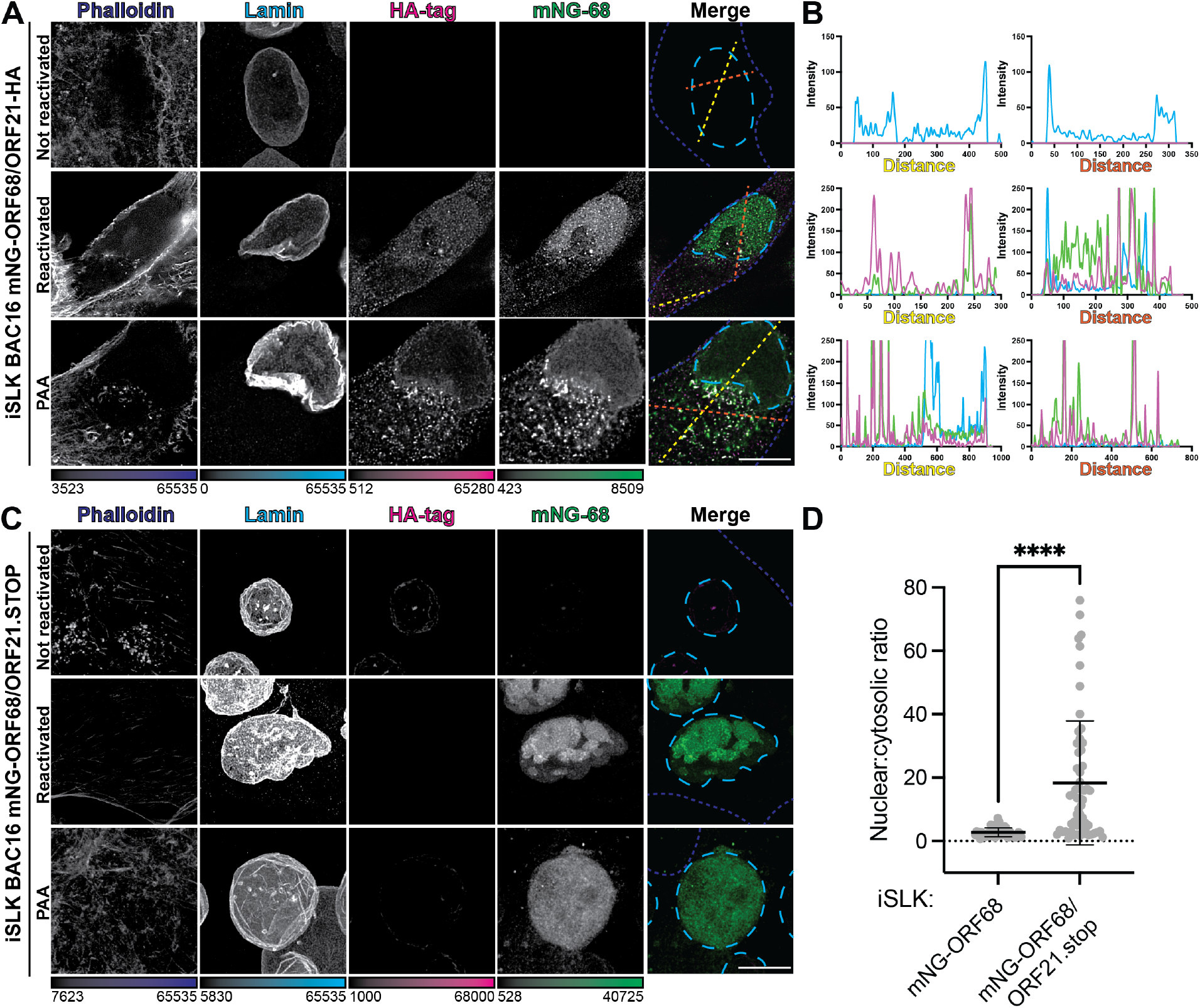
ORF68 localization to cytoplasmic puncta is contingent on expression of ORF21. **A**. Z-slices of iSLK mNG-ORF68/ORF21-HA cells. Cells were fixed 72 hours post-reactivation, with or without PAA treatment, imaged for mNG-ORF68 (green), and stained for phalloidin (blue dashed lines), lamins A and C (cyan dashed lines), and HA-tag (magenta). Representative of at least 10 cells from two biological replicates. **B**. Intensity profile plots showing intensity of Lamin (blue), mNG (green), or HA-tag (magenta) signal at each point along lines (yellow or orange dashes). **C**. Whole-cell maximum Z-projections of iSLK mNG-ORF68/ORF21.stop cells. Cells were imaged for mNG-ORF68 (green) and stained for phalloidin (blue dashed lines), lamins A and C (cyan dashed lines), and HA-tag (magenta), which should be undetectable in cells lacking an HA epitope tag. Representative of at least 10 cells from two biological replicates. Scale bars are 10 μm; color bars represent the modified minimum and maximum pixel intensity values. **D**. Quantification of nuclear to cytosolic ratio of mNG-ORF68 in the presence or absence of viral-expressed ORF21. Error bars indicate the mean, S.D. (***) indicates a p-value < 0.0001 calculated using Welch’s T-test.

Next, we asked if localization of ORF68 to cytoplasmic puncta was contingent on expression of ORF21. We introduced a premature termination codon in the ORF21 coding region to prevent ORF21 protein expression (ORF21.stop) in the mNG-ORF68 BAC and generated latently infected iSLK cell lines (**Fig. S4A**). ORF21 has previously been reported to be nonessential for virion production in iSLKs^70^, and indeed these cell lines were generated in the absence of complementing ORF21 expression. Expression of ORF68 and other representative viral genes was unchanged in the absence of ORF21 (**Fig. S4B**). We stained for lamins A/C and phalloidin to demarcate the nucleus and cytoplasm and monitored ORF68 localization via its mNG tag. Strikingly, in the absence of ORF21, ORF68 fails to form cytoplasmic puncta, and its localization was predominantly nuclear (**Fig. 4C, D**). Interestingly, ORF68 localizes to the nucleus upon PAA treatment, suggesting that it is recruited and retained in the nucleus even in the absence of capsids.

### The disordered N-terminal domain of ORF21 controls ORF68 subcellular localization

Finally, to establish whether ORF68 and ORF21 interact in the absence of other viral factors, we monitored ORF68 localization outside the context of infection. We generated plasmids encoding ORF68 fused to an N-terminal HA tag and ORF21 fused to a C-terminal 2xStrep tag, then transiently transfected HEK293T cells and visualized by IF. In the absence of other viral factors, ORF68 localizes primarily to the nucleus, with limited cytoplasmic localization (**Fig. 5A**). This morphology is consistent with prior observations of HSV-1 homolog UL32 localization upon transient transfection^39^. In contrast, full-length ORF21 is exclusively cytoplasmic (**Fig. 5A**). When ORF21 is co-transfected with ORF68, both proteins exhibit cytoplasmic localization, suggesting that an interaction with ORF21 is sufficient to prevent ORF68 nuclear localization.

**FIGURE 5.**
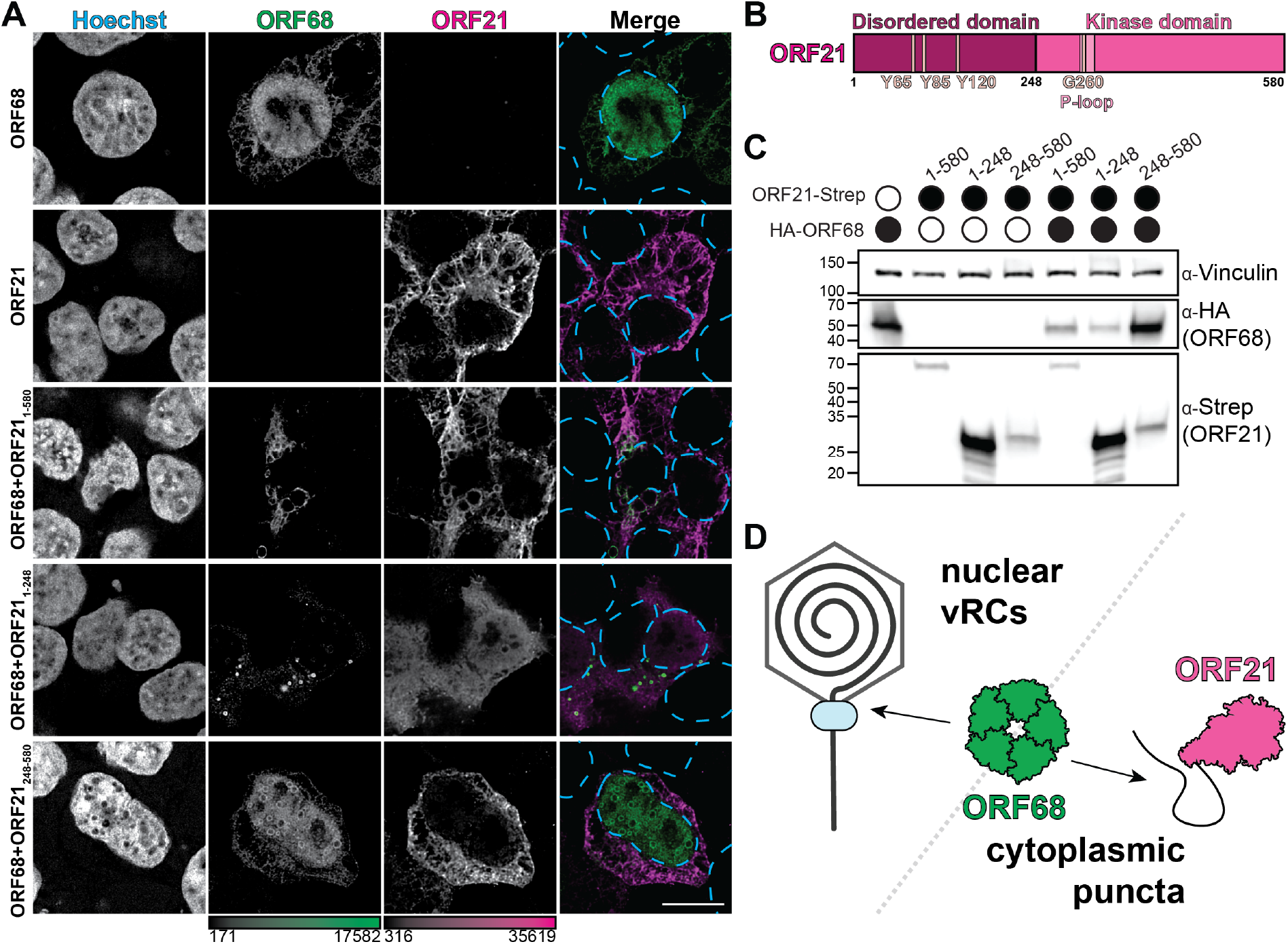
The N-terminus of ORF21 drives cytoplasmic sequestration of ORF68. **A**. Z-slices of transfected HEK293T cells expressing HA-ORF68 and/or ORF21-Strep constructs, including ORF21 full-length, N-terminal region (residues 1-248), and kinase domain (residues 248-580). Stained for Hoechst (cyan dashed lines), HA-tag (green), and Strep-tag (magenta), representative of at least 10 cells from 2 biological replicates. Color bars represent the modified minimum and maximum pixel values. Scale bar is 10 μm. **B**. Schematic of ORF21 domain organization, showing disordered region and kinase domain. **C**. Western blot of whole cell lysate (25 μg) from HEK293T cells transfected with ORF21 constructs (Strep) and ORF68 (HA). Vinculin serves as the loading control. **D**. ORF68 participates in viral genome packaging in nuclear viral replication compartments, but is drawn to the cytoplasm through an interaction with the disordered N-terminal tail of ORF21.

ORF21 consists of an unstructured N-terminal domain (residues 1-248) and a highly structured C-terminal kinase domain (248-580) that closely resembles the canonical thymidine kinase fold including a catalytic P-loop motif^69^ (**Fig. 5B**). However, ORF21 is an inefficient thymidine kinase with weak affinity for thymidine^72^. Instead, ORF21 is a tyrosine kinase that autophosphorylates three tyrosine residues in its N-terminal domain (Y65, Y85, and Y120) and influences cell morphology during infection^69^. We tested whether the N-terminal or kinase domains were responsible for influencing ORF68 localization. When the ORF21 kinase domain is co-expressed with ORF68, ORF68 localizes to the nucleus and cytoplasm, while ORF21_248-580_ resides in the cytoplasm, similar to when the full-length proteins are expressed separately (**Fig. 5A**). In contrast, while the ORF21 N-terminal domain localized to both the nucleus and cytoplasm, its expression restricted ORF68 to the cytoplasm of transfected cells and induced ORF68 puncta (**Fig. 5A**). To test if the cytoplasmic colocalization of ORF68 and ORF21 is dependent on autophosphorylation of the N-terminal tyrosine residues in ORF21, we co-transfected ORF68 with full-length ORF21 bearing an alanine mutation at position Y120. We also introduced a mutation to the kinase active site (G260V) predicted to reduce ATP binding^73^. This mutation did not restore nuclear ORF68 localization, suggesting that the ORF68-ORF21 interaction is not dependent on ORF21 kinase activity or autophosphorylation at ORF21 Y120 (**Fig. S5**). Interestingly, expression of full-length or N-terminal ORF21 reduced ORF68 expression by both IF and western blot (**Fig. 5C**). However, we did not observe an increase in ORF68 expression upon deletion of ORF21 in iSLK cells (**Fig. S4B**). We propose that ORF21 controls the abundance of nuclear ORF68 available for viral genome packaging, potentially serving as a lever by which to regulate virion production (**Fig. 5D**).

## DISCUSSION

All herpesviruses replicate their genomes and assemble capsids in the nuclei of infected cells. We find that during KSHV lytic replication, the packaging accessory factor ORF68 localizes to both nuclear replication compartments and previously uncharacterized cytoplasmic puncta. Using proximity biotinylation, we identified an interaction between ORF68 and ORF43, the capsid portal protein, suggesting that the packaging accessory factor is physically proximal to the capsid during viral genome packaging. Three lines of evidence suggest that ORF68 may serve an additional function independent of its canonical role in packaging. First, cytoplasmic ORF68 puncta can be observed as early as 24 hours post-reactivation, before infectious virions are produced in the iSLK cell culture model. Second, we observed these puncta upon PAA treatment, an inhibitor of viral genome replication that prevents late gene expression and thus assembly of nascent capsids for packaging. Third, we show that other components of the packaging machinery and capsid do not colocalize with cytoplasmic ORF68 puncta.

Localization of ORF68 to cytoplasmic puncta is dependent on the viral tyrosine kinase ORF21. ORF68 and its homologs are conserved across the *Herpesviridae* and required for viral genome packaging in all models tested^29,39,41^. In contrast, ORF21 is poorly conserved and lacks a homolog in the betaherpesvirus subfamily. Gammaherpesvirus ORF21 homologs consist of a kinase domain with a long N-terminal intrinsically disordered domain. Alphaherpesviruses encode true viral thymidine kinases (i.e., HSV-1 UL23), where the structurally conserved kinase domain is preceded by a shorter N-terminal extension. Interestingly, IF of the ORF68 homolog in HSV-1 revealed that it forms cytoplasmic puncta^74^, while imaging of the homolog from the betaherpesvirus HCMV revealed exclusively diffuse nuclear localization^43^. Thus, the viral kinase in alpha- and gammaherpesviruses may affect localization of the packaging accessory factor, with differences in the length or identity of the intrinsically disordered N-terminal region dictating distinct localization phenotypes.

ORF21 could potentially modulate ORF68 activity through installation of phosphotyrosine residues. Furthermore, the subcellular localization of ORF68 to the vRC or the cytoplasmic puncta could also be mediated by its phosphorylation state, which could be altered by ORF21 activity. Phosphoproteomic analysis of viral proteins during murine gammaherpesvirus (MHV68) infection indicated that the murine ORF21 homolog is post-translationally modified during infection, including several phosphotyrosines^75^. However, only a single phosphoserine was detected on the murine ORF68 homolog. Despite the close evolutionary relationship of KSHV and MHV68, ORF21 appears to exhibit different functional properties in the two viruses^69,71^. ORF21 is nonessential in the KSHV iSLK cell culture model, allowing us to disentangle the essential nuclear packaging role of ORF68 and secondary cytoplasmic role through use of the iSLK ORF21.stop cell line. However, ORF21 is required for gammaherpesvirus infection in mice, indicating that ORF21 plays a key role during pathogenesis^76,77^. If the ORF68-ORF21 interaction is recapitulated in MHV68, future work could use a murine model to investigate the significance of these novel cytoplasmic puncta.

One of the top hits in our proximity labeling dataset was the phosphoinositide 3-kinase 1-alpha (PI3K1α) complex, which is a heterodimer composed of the p110 catalytic subunit (PIK3CA) and p85 subunit (PIK3R1.1). Other PI3K family subunits (PIK3R2, PIKFYVE, PIK3R1, PIK3R3, PIK3CB, PIK3CB.1, PIK3R1.2, PIK3C2B, PIK3C2A, PIK3CD, PIK3C3, and PIK3R4) were detected but not enriched above the controls, allowing for specific identification of PI3K1α. ORF21 has previously been shown to interact with p85 outside the context of infection^69^, suggesting that PI3K1α is detected in our ORF68 proximity labeling dataset by virtue of its interaction with ORF21. Herpesviruses modulate the PI3K1α pathway during infection, leading to changes in autophagy, metabolism, and other homeostatic processes^75,78–81^. We speculate that the cytoplasmic function of ORF68 and ORF21 is to modulate the PI3K1α pathway, resulting in pro-viral changes to host cell autophagy and apoptosis. Other KSHV proteins – vBcl-2^82^, vFLIP^83^, and K7^84^ – are known to alter autophagy. Like ORF21, the anti-autophagy/apoptosis function of these proteins are not essential for KSHV virion production *in vitro*, although these functions are essential for viral success in more complex models. Unlike in other herpesviruses, both ORF68 and ORF21 are detected in mature KSHV virions^85^. Whether recruitment of ORF68 to the KSHV tegument depends on ORF21 remains to be determined, and whether delivery of ORF68/ORF21 to a new cell influences its fate.

ORF68 forms a pentameric ring^40^. We hypothesize that further oligomerization into a biomolecular condensate occurs through multivalent interactions between ORF68 pentamers and the N-terminal intrinsically disordered region of ORF21, forming discrete puncta in the cytoplasm of infected cells. Due to the comparatively small size of herpesvirus exomes, many herpesvirus proteins have evolved to fulfill multiple roles throughout the course of lytic infection. Our work reveals a potential novel secondary function of a conserved packaging factor, highlighting how species-specific differences in protein-protein interactions can drive unique biology.

## ACKNOWLEDGEMENTS

We thank Dr. Joe Wolenski and Dr. Binyam Mogessie for advice on microscopy data collection and analysis. We thank Dr. Tom Melia and Dr. Mandy Muller for advice and feedback. We are thankful to all members of the Didychuk lab for feedback. We thank the Yale University Light Microscope Imaging Facility on Science Hill for usage of their Leica STELLARIS 8 TauSTED confocal microscope.

## MATERIALS AND METHODS

### Plasmids

The vector for transient expression of ORF21 was previously described (pcDNA4/TO-ORF21-2xStrep, Addgene #136181)^86^. Mutations and truncations to ORF21 (ORF21 Y120A, G260V, 1-248, and 258-580) were introduced by inverse PCR site-directed mutagenesis using primers listed in **Supplementary Table S2**. PCR products were treated with DpnI for 1 hour at 37°C then ligated using T4 DNA ligase and T4 PNK (New England Biolabs) for 2 hours at room temperature before transformation into *E. coli* 5-alpha competent cells (New England Biolabs). Plasmids were confirmed by Sanger sequence and deposited to Addgene (#252281-252285). The HA-ORF68 transient expression plasmid (Addgene #252278) was constructed using inverse PCR as above using pcDNA4/TO-2xStrep-ORF68 (Addgene #162625) as a template.

### BAC mutagenesis

Red recombineering of KSHV BAC16 was performed using a double-stranded DNA “geneblock” harboring a kanamycin resistance cassette flanked by homologous regions containing the mutation or insertion of interest^87^. We used geneblocks where the right homology arm consisted of 125 bp upstream of the mutation, the mutation/insertion, and 25 bp downstream of the mutation. The left homology arm consisted of 25 bp upstream of the mutation, the mutation/insertion, and 125 bp downstream of the mutation. Geneblocks (**Supplementary Table S3**) were synthesized by Integrated DNA Technologies or amplified from plasmids, which we generated by amplifying homology arms from synthetic gene fragments (Twist Biosciences) or the BAC16 genome, and using In-Fusion cloning (Takara Bio) to ligate along with the kanamycin cassette into pUC119 digested with HindIII and EcoRI (New England Biolabs).

Scarless Red recombination mutagenesis of KSHV BAC16 was performed as previously described^87^. Briefly, GS1783 *E. coli* containing BAC16 were electroporated with double-stranded DNA harboring a kanamycin resistance cassette flanked by homologous regions containing the mutation or insertion of interest (“geneblocks”) (**Supplementary Table S3**). Successful recombinants were selected on CAM/KAN LB-agar plates for 2 days at 30°C. A second recombination to remove the kanamycin resistance cassette was performed in the presence of 1% arabinose. Selection occurred on CAM/1% arabinose LB-agar plates. Successful mutagenesis was confirmed by Sanger sequencing of the locus. BAC DNA was purified using a Nucleobond Xtra BAC kit (Macherey-Nagel) and integrity of the BAC was assessed through restriction digestion with RsrII (New England Biolabs) and analysis on a 0.75% agarose gel with SybrSafe staining.

### Mammalian cell culture

HEK293T cells (ATCC CRL-3216) were cultured at 37°C with 5% CO_2_ in Dulbecco’s modified eagle media (DMEM) supplemented with 10% fetal bovine serum (FBS). iSLK-puro cells were cultured in DMEM+10% FBS with 1 μg/mL puromycin. After establishment of latent infection with KSHV encoded on BAC16, iSLK-BAC16 cells were cultured in DMEM+10% FBS with 1 μg/mL puromycin and 1 mg/mL hygromycin.

### Establishment of KSHV-infected cell lines

HEK293T cells were transfected with 10 μg of BAC DNA using 30 μg PEImax MW 40,000 (Polysciences, Inc). The next day, 1 million transfected HEK293T cells were cultured with 1 million iSLK-puro cells. After 4 hours, cells were induced with 25 nM 12-O-Tetradecanoylphorbol-13-acetate and 0.3 mM sodium butyrate. Three days later, fresh DMEM + FBS was added. The next day, cells were trypsinized and selected with 1 μg/mL puromycin, 300 μg/mL hygromycin, and 250 μg/mL G418. After one week of selection, the hygromycin concentration was progressively increased to a final selection concentration of 1000 μg/mL.

### TurboID biotinylation and enrichment

Biotinylation and affinity purification were performed as previous described^88^. Two 15-cm plates were seeded with 10 million iSLK cells. The next day, cells were reactivated with 5 μg/mL doxycycline and 1 mM sodium butyrate for 48 hours. Cells were washed with DPBS and fresh media supplemented with 500 μM biotin (MilliporeSigma) was added followed by incubation at 37°C for 15 minutes. To allow for incorporation of any free biotin, the biotin-containing media was replaced by fresh media and incubated at 37°C for an additional 30 minutes. Cells were harvested in cold DPBS, pelleted at 300 x g for 5 minutes at 4°C and frozen at -80°C. All three replicates for the samples were processed in parallel. Cell pellets were resuspended in RIPA buffer (50 mM TRIS pH 8, 150 mM NaCl, 0.1% SDS, 1% Triton X-100, 0.5% sodium deoxycholate) supplemented with protease inhibitors (cOmplete protease inhibitors EDTA-free, Roche) and rotated at 4°C for 1 hour. Samples were treated with benzonase for 15 minutes at 37°C then sonicated at 100 amps in a cuphorn water-bath (QSonica) for a total time of 1 min (3 seconds on, 17 seconds off) at 4°C. Lysates were clarified by centrifugation at ∼12,000 x g for 10 minutes at 4°C. Total protein concentration in the supernatant was measured using a Bicinchoninic acid (BCA) protein assay (Pierce).

Biotinylated proteins were enriched through affinity purification with 50 μL magnetic streptavidin beads (Pierce) and 6 mg of total protein in a final volume of 6 mL in RIPA buffer containing protease inhibitors (Roche). Samples were rotated for 1 hour at room temperature and then overnight at 4°C. The beads were washed for the indicated times with 1 mL of RIPA buffer (2 minutes, twice), 1M KCl (2 minutes, once), 100 mM Na_2_CO_3_ (2 minutes, once), 2M Urea in 10 mM Tris-HCl pH 8 (10 seconds, once), and RIPA buffer (2 minutes, twice). Beads were stored in RIPA buffer at 4°C for prior to processing for mass spectrometric analysis. For western blotting analysis, the beads were washed three times with 1 mL wash buffer (50 mM TRIS pH 7.4, 150 mM NaCl) prior to elution by boiling in 1x Laemelli buffer (Bio-Rad) supplemented with 100 mM biotin.

### TurboID sample preparation for mass spectrometry

Affinity purified samples on streptavidin beads were transferred to a new Low Bind tube (Amuza, Inc.), and storage RIPA buffer was removed. Beads were washed twice with 200 µL 50 mM Tris HCl pH 7.5 and twice with 1 mL 2 M urea in 50 mM Tris pH 7.5. An on-bead digestion of the 4 mg sample was performed by resuspending the beads in 80 µL 2 M urea in 50 mM Tris pH 7.5 with 1 mM DTT and 3.2 µg trypsin and incubating for 1 h at 25°C while shaking at 600 rpm. Supernatant was transferred to a new Low Bind tube, the beads were washed twice with 60 uL 2 M urea in 50 mM Tris pH 7.5, and the washes were combined with the on-bead digest supernatant. Samples were reduced and alkylated with 25 mM TCEP (Tris(2-carboxyethyl)phosphine) (ThermoFisher) and 50 mM CAM (chloroacetamide) (Fisher Scientific) by shaking at 600 rpm in a thermomixer at 25°C for 1.5 h. Samples were diluted 1:1 (v/v) with 50 mM Tris pH 8 and an additional 4 µg trypsin was added. Samples were digested overnight at 25°C with shaking at 600 rpm. Samples were acidified to 1% trifluoroacetic acid, incubated on ice for 15 min, and spun down at 4000 x g for 5 min at 4°C. Samples were desalted over C18 StageTips (Empore™ SPE Disks C18, Fisher Scientific) and dried in a speedvac before resuspension in 6 µL 1% formic acid/1% acetonitrile.

### Mass spectrometry data acquisition

Samples were analyzed on a timsTOF Ultra mass spectrometer (Bruker) equipped with a Captive Spray 2 ion source (Bruker) containing a 10 μm emitter (Bruker). Peptides were resolved for nLC-MS/MS using a nanoElute 2 nLC system (Bruker) equipped with a PepSep C18 column (1.5 μm by 75 μm by 25 cm; Bruker). Peptides were separated with a 40-min gradient using a mobile phase composed of 0.1% FA as solvent A and 0.1% FA/99.9% ACN as solvent B. A linear gradient was run consisting of 3 to 34% buffer B at a flow rate of 200 nl/min. Mass spectrometry data were acquired with TIMS enabled in dia-PASEF mode. The instrument was operated in positive ion polarity over an m/z range of 100–1700. TIMS settings were configured in Custom mode with 1/K0 start = 0.65 V·s·cm^−2^ and 1/K0 end = 1.55 V·s·cm^−2^, using a ramp time of 50.0 ms and an accumulation time of 50.0 ms ( duty cycle = 100.00%; ramp rate = 17.80 Hz ). In-batch calibration was set to OFF. The ion source was operated in ESI mode with end plate offset = 500 V, capillary voltage = 4500 V, nebulizer = 0.2 bar, dry gas = 3.0 L/min, and dry temperature = 180 °C. Tune settings were as follows. Transfer: deflection 0 delta 110.0 V, funnel 0 RF 250.0 Vpp, multipole 0 RF 100.0 Vpp, deflection 1 delta 70.0 V, funnel 1 RF 450.0 Vpp, isCID energy 0.0 eV, funnel 2 RF 200.0 Vpp, and multipole RF 500.0 Vpp. Quadrupole: ion energy 5.0 eV and low mass 200.00 m/z. Focus Pre-TOF: transfer time 60.0 µs and pre-pulse storage 12.0 µs. Collision cell: collision energy 10.0 eV and collision RF 1500.0 Vpp. The High Sensitivity Detection – Low Sample Amount option was not enabled.

### Mass spectrometry data analysis

Data were analyzed using DIA-NN 1.8 (91). MS/MS spectra were searched using a FASTA containing human (downloaded January 2021) and KSHV protein sequences (downloaded March 2021) and common contaminants using trypsin as the digestion enzyme. Included modifications were static cysteine carbamidomethylation, dynamic methionine oxidation, dynamic N-terminal acetylation, and dynamic methionine excision at the N terminus. A maximum of one missed cleavage and one variable modification were allowed. Default settings were used for both searches. The mass accuracy was set at 10 ppm, the MS1 accuracy at 7 ppm, and a scan window of 10. “Match between runs” was enabled. The proteomics output tables were filtered for a maximum of 1% q value at both precursor and global protein levels. Two peptides per protein were required, with only unique or razor peptides used for quantification. Differential expression analysis was performed using Fragpipe analyst^89^. Subcellular localization was assigned based on Uniprot annotations. Protein networks were generated using Cytoscape (v.3.8.2)^90^ and functional module detection was generated using Humanbase^91^.

### Supernatant transfer assay

10-cm plates were seeded with 1 million iSLK BAC16 cells. The next day, cells were reactivated with 1 mM doxycycline and 5 μg/mL sodium butyrate and incubated at 37°C for 72 hours. Virus-containing supernatants were 0.45 μm filtered and spinfected (870 x g for 2 hours at 37°C) onto 1 million HEK293T cells in a 6-well plate, then incubated at 37°C for 24 hours. The HEK293T cells were washed 3 times with DPBS and fixed in 4% PFA in DPBS for 10 minutes, rotating at 4°C. Then, cells were washed with DPBS, stored in DPBS, filtered through 35 μm cell strainers (Falcon), and analyzed by flow cytometry.

### Western blot

10-cm plates were seeded with 1 million iSLK BAC16 cells. The next day, cells were reactivated with 1 mM doxycycline and 5 μg/mL sodium butyrate and incubated at 37°C for 72 hours. Cell pellets were resuspended in lysis buffer (50 mM TRIS pH 7.4, 150 mM NaCl, 1 mM EDTA, 0.5% Nonidet P-40, cOmplete protease inhibitor) and rotated for 1 hour at 4°C. Samples were centrifuged at ∼12,000 x g for 10 minutes at 4°C and the supernatant was decanted. The total protein concentration of the supernatant was assessed with a Bradford assay. Samples for SDS-PAGE were prepared in 2X Laemmli Buffer, run on a 4-20% Mini-PROTEAN TGX SDS-PAGE gel (Bio-Rad) and transferred to PVDF nitrocellulose membrane. The membrane was blocked in 5% milk in TBST (Tris-buffered saline with 0.2% Tween-20) at room temperature, then incubated in primary antibody (**Supplementary Table S4**) in 0.5% milk overnight at 4°C. Then, the membrane was washed three times with TBST and incubated in secondary antibody (**Supplementary Table S4**) for 1 hour at room temperature prior to visualization with Clarity Western ECl (Bio-Rad) on an Azure 600 imager.

### Viral replication qPCR assay

6-well plates were seeded with 1 million iSLK BAC16 cells per well. The next day, cells were either left unreactivated or reactivated with 1 mM doxycycline and 5 μg/mL sodium butyrate. Cells were incubated for 72 hours at 37°C. Subsequently, cells were washed with ice-cold DPBS and pellets were harvested and stored at -80°C prior to DNA isolation with a NucleoSpin Blood kit (Macherey Nagel). qPCR was performed with iTaq UniverSYBR Green (Bio-Rad) on reactivated (R) and non-reactivated (NR) samples, with primer pairs against the promoters of KSHV ORF57 (ORF57Pr) or human CTGF (CTGFPr) (**Supplementary Table S2**). For each sample, fold change was calculated using the ΔΔC_T_ method (2^(- ([C_T(R, ORF57Pr)_- C_T(R, CTGFPr)_]- [C_T(NR, ORF57Pr)_- C_T(NR, CTGFPr)_]))).

### Virion release qPCR

6-well plates were seeded with 1 million iSLK BAC16 cells per well. The next day, cells were either left unreactivated or reactivated with 1 mM doxycycline and 5 μg/mL sodium butyrate, and optionally treated with PAA. Cells were incubated for 72 hours at 37°C. Virus-containing supernatants were 0.45 μm filtered, and subsequently treated with 30 U DNaseI (ThermoFisher) for 1 hour at 37°C. DNase was heat inactivated for 10 minutes at 75°C. Samples were treated with 400 μg/mL Proteinase K (GoldBio) in 0.5% SDS overnight at 37°C, then heat inactivated for 30 minutes at 65°C. Samples were prepared with a NucleoSpin Blood kit (Macherey Nagel).

qPCR was performed with iTaq UniverSYBR Green (Bio-Rad) on reactivated (R), non-reactivated (NR), or phosphonoacetic acid (PAA)-treated samples, with primer pairs against the promoters of KSHV ORF57 (ORF57Pr) or human CTGF (CTGFPr) (**Supplementary Table S2**). A standard curve for the KSHV genome was produced using a 6-point serial dilution of purified KSHV BAC16, to convert KSHV concentration to C_T,ORF57Pr_ value. For each sample, C_T,ORF57Pr_ was converted to concentration and then subsequently KSHV genome copy number using the molecular weight of the genome.

### Immunofluorescence of iSLK cells

100-200K iSLK BAC16 cells were seeded onto sterile No. 1.5 coverslips. 24 hours later, cells were reactivated with 1 mM doxycycline and 5 μg/mL sodium butyrate. Cells were incubated for 72 hours at 37°C. Then, cells washed 3 times with DPBS and subsequently fixed in 4% paraformaldehyde (PFA) for 10 minutes. Cells were permeabilized with 0.5% Triton X-100 for 10 minutes. Samples were blocked with 3% Bovine Serum Albumin in DPBS for 30 minutes (Sigma). Samples were incubated with primary antibodies overnight at 4°C (**Table S2)**. Subsequently, samples were incubated with secondary antibodies for 1 hour at room temperature. Samples were optionally incubated in 1:2000 Hoechst 33342 (Invitrogen) for 20 minutes. Coverslips were mounted on slides with Diamond ProLong Antifade Mountant (Invitrogen) and cured for 24 hours.

### EdU labeling of iSLK cells

100-200K iSLK BAC16 cells were seeded onto sterile No. 1.5 coverslips. 24 hours later, cells were reactivated with 1 mM doxycycline and 5 μg/mL sodium butyrate for 70 hours at 37°C prior to EdU labeling using the Click-iT™ Plus EdU Cell Proliferation Kit for Imaging, Alexa Fluor™ 594 dye (Invitrogen) following the manufacturer’s protocol. Briefly, cells were incubated with 10 μM EdU in DMEM supplemented with 10% FBS for 2 hours at 37°C prior to fixation in 4% PFA in DPBS for 15 minutes. Cells were washed twice in DPBS, then twice in 3% BSA in DPBS. Cells were perforated in 0.5% Triton X-100 in DPBS for 20 minutes at room temperature. Then, cells were treated with Click-iT Plus reaction cocktail for 30 minutes at room temperature. Samples were incubated in 1:2000 Hoechst (Invitrogen) for 20 minutes. Coverslips were mounted on slides with Diamond ProLong Antifade Mountant (Invitrogen) and cured for 24 hours.

### Transient transfection and immunofluorescence of HEK293T cells

100-200K HEK293T cells were seeded onto sterile No. 1.5 coverslips. 24 hours later, cells were transiently transfected with 2 μg DNA using 5 μg PEImax MW 40,000 (Polysciences, Inc). 24 hours later, cells were washed 3 times with DPBS and fixed in 4% PFA in DPBS for 10 minutes. Cells were permeabilized with 0.5% Triton X-100 for 10 minutes. Samples were blocked with 3% Bovine Serum Albumin (Sigma) in DPBS for 30 minutes, then incubated with primary antibodies overnight at 4°C (**Table S2)**. The next day, samples were incubated with secondary antibodies for 1 hour at room temperature. Samples were incubated in 1:2000 Hoechst (Invitrogen) for 20 minutes. Coverslips were mounted on slides with Diamond ProLong Antifade Mountant (Invitrogen) and cured for 24 hours.

### Microscopy

Imaging was performed on a Leica STELLARIS 8 TauSTED confocal microscope outfitted with a 405nm DMOD laser and a white light laser (440 nm to 790 nm) with five Power Hybrid detectors (HyD) using a Leica HC PL APO 100x/1.40 oil objective with Immersion Oil 518F (Zeiss). Images were captured in 2 sequences (stack sequential) to avoid crosstalk. Deconvolution was performed using LIGHTNING (Leica). Processing was performed in LAS X and Fiji (ImageJ v1.54p)^92^. Intensity profile plots were generated in Fiji using the RGBProfiles macro (https://imagej.net/ij/macros/tools/RGBProfilesTool). Quantitative analysis of nuclear vs. cytoplasmic mNG-ORF68 was performed by measuring the average intensity of mNeonGreen in a square with an area of 14.5 nm in the nucleus (determined using lamin A/C staining) and in the cytoplasm (determined using phalloidin staining). Images of a single Z-stack where both the nucleus and cytosol were in focus were analyzed using Fiji. Subsequently, the ratio of nuclear to cytosolic signal was determined and outliers were removed using ROUT method with Q = 0.5%. Welch’s T-test was used to determine statistical significance.

## Data Availability

The mass spectrometry proteomics data have been deposited to the ProteomeXchange^93^ Consortium via the PRIDE^94^ partner repository with the dataset identifier PXD073504.

---

**SUPPLEMENTARY FIGURE S1.**
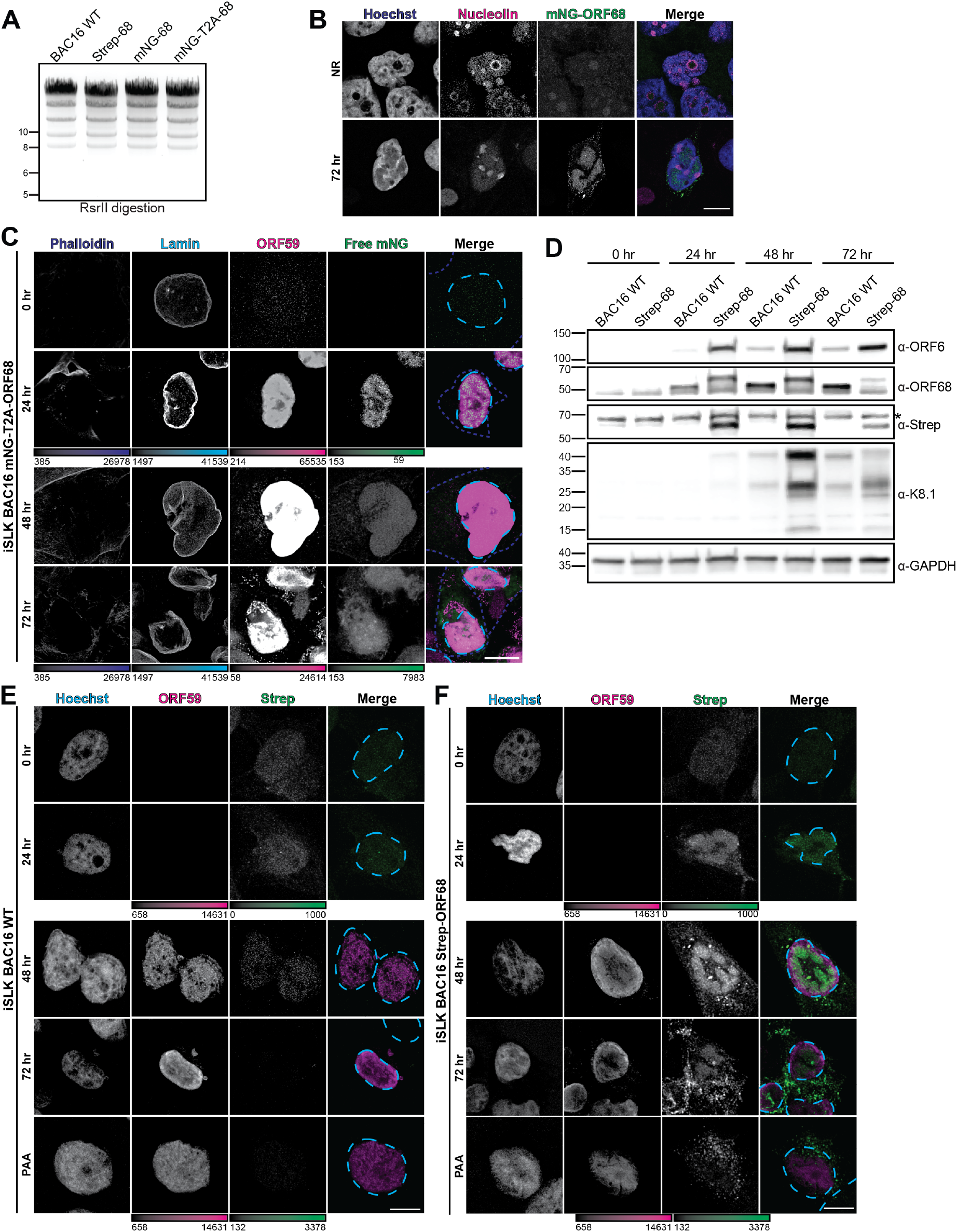
Alternate tagging of ORF68 produces a similar punctate phenotype unrelated to mNG expression. **A**. Recombinant TSP-ORF68, mNG-ORF68, and mNG-T2A-ORF68 BACs were digested with RsrII to assess that no large-scale rearrangements occurred during cloning. **B**. Z-slices of iSLK mNG-ORF68 cells at 72 hours post-reactivation, stained for Hoechst (blue) and nucleolin (magenta), representative of at least 5 cells from 2 biological replicates. **C**. Z-slices of iSLK cells expressing mNG-T2A (green) separately from ORF68 at 0, 24, 48, and 72 hours post reactivation. Stained for phalloidin (blue dashes), Lamin A/C (cyan dashes), and ORF59 (magenta), representative of at least 5 cells from 2 biological replicates. **D**. Western blot of whole cell lysate (25 μg) from iSLK WT or TSP-ORF68 cells at 0, 24, 48, or 72 hours post-reactivation. We blotted for levels of ORF68, early genes (ORF6), and late genes (K8.1, ORF26). GAPDH is the loading control. Asterisk (*) indicates a nonspecific band. 3 biological replicates. **E**. Z-slices of lytic iSLKs expressing WT ORF68 at 0, 24, 48, and 72 hours post reactivation, with or without PAA treatment. Stained for Hoechst (cyan dashed line), Strep (green), and ORF59 (magenta), representative of at least 5 cells from 1 biological replicate. **F**. Z-slices of lytic iSLK TSP-ORF68 cells at 0, 24, 48, and 72 hours post reactivation, with or without PAA treatment. Cells were stained for Hoechst (cyan dashed line), Strep (green), and ORF59 (magenta), representative of at least 5 cells from 1 biological replicate. Scale bars are 10 μm; color bars represent the modified minimum and maximum pixel values.

**SUPPLEMENTAL FIGURE S2.**
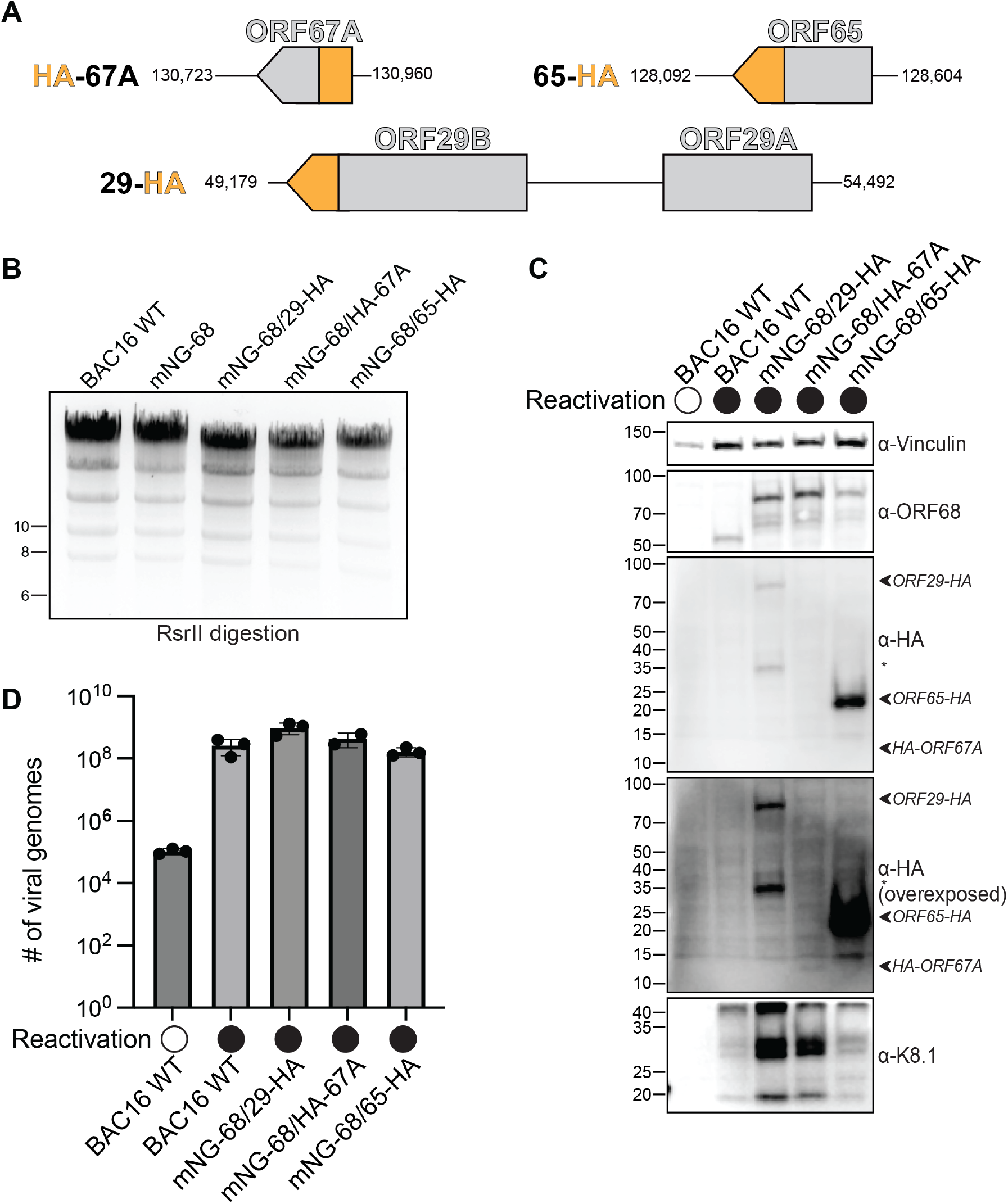
Tagging ORF68 and capsid or packaging proteins simultaneously does not disrupt the lytic cycle. **A**. Schematic of the loci in the BAC16 WT genome and insertion of HA-tag at the N-terminus of ORF67 (HA-67), C-terminus of ORF65 (65-HA), and C-terminus of ORF29 (29-HA). **B**. Recombinant mNG-ORF68, mNG-ORF68/ORF29-HA, mNG-ORF68/HA-ORF67A, and mNG-ORF68/ORF65-HA BACs were digested with RsrII to assess that no large-scale rearrangements occurred during cloning. **C**. Western blot of whole cell lysate from reactivated iSLK cells indicate that HA-tagged ORFs (labeled with arrows) are detectable by western blot and tagging does not influence expression of representative early (ORF68) or late (K8.1) genes, representative of three biological replicates. The asterisk refers to an isoform of ORF29-HA. Vinculin is the loading control. **D**. Extracellular virion production of unreactivated or reactivated iSLK cells was quantified by qPCR from three biological replicates. Filled circles indicate the number of viral genome copies measured for each biological replicate, with the bar representing the mean and error bars depicting the SD.

**SUPPLEMENTARY FIGURE S3.**
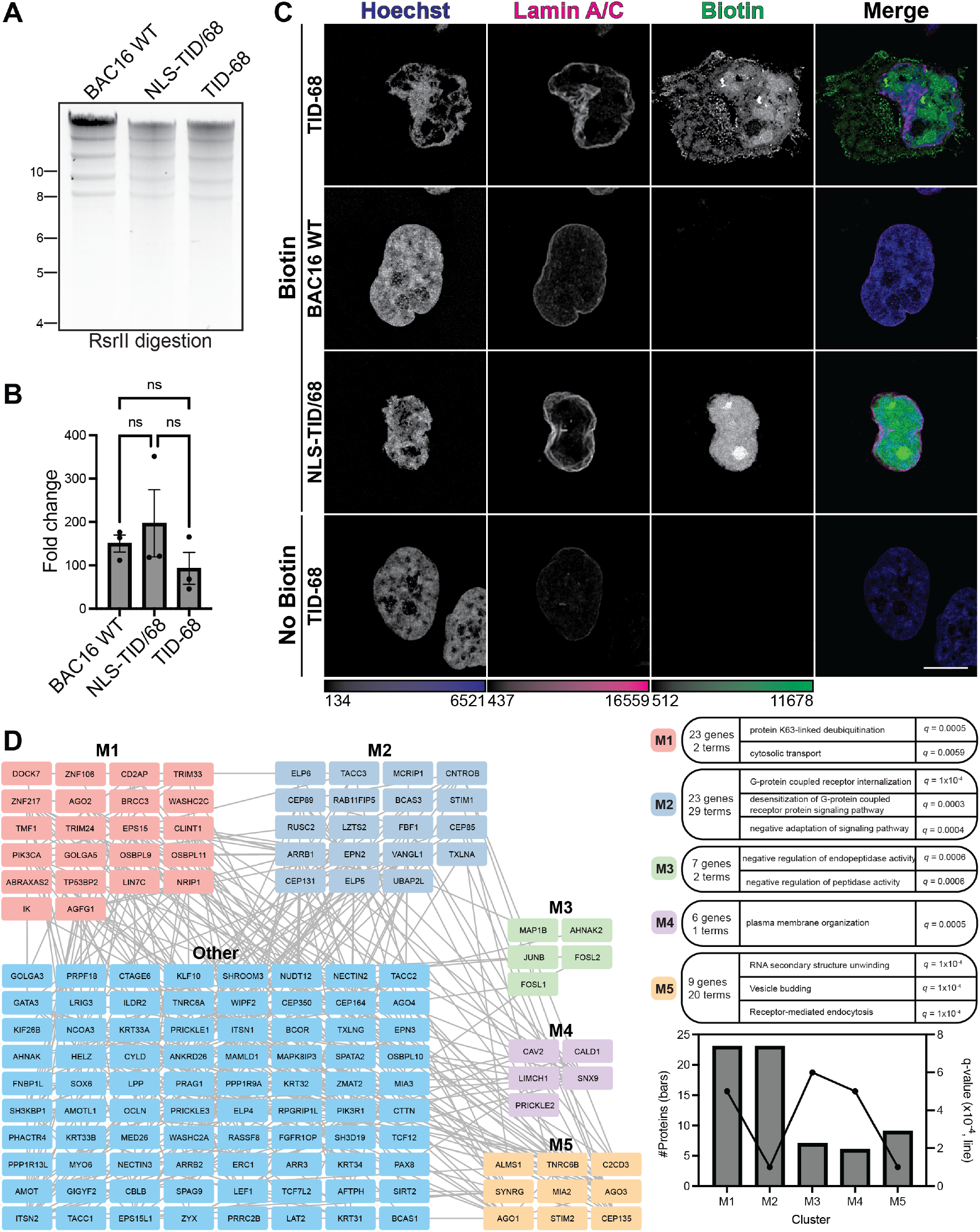
**A**. RsrII BAC digestion of BAC16 WT compared to engineered BACS containing NLS-TID/68 and TID-68. **B**. qPCR to quantify viral genome replication at 72 hours post-reactivation of iSLKs infected with BACs WT, NLS-TID/68, or TID-68. Genome replication is not significantly (ns) different between the viruses by RM one-way ANOVA (p>0.05). 3 biological replicates. **C**. Z-slices of lytic iSLKs expressing ORF68 WT (BAC16 WT), TurboID-ORF68 (TID-68), or TurboID-T2A-68 (NLS-TID/68) at 48 hours post reactivation, fixed after 15-minute biotin (“Biotin”) or DMSO (“No Biotin”) treatment. Stained for Hoechst (blue) and Lamin A/C (magenta). Streptavidin-conjugated to a fluorophore demarcates biotinylated proteins (green). Representative of at least 5 cells from one biological replicate. Scale bar is 10 um. **D**. Proximity-based interaction network of ORF68-specific host proteins. GO term clusters were generated with HumanBase using functional module detection^97^. Bar and line plot indicates the number of proteins and q-values associated with each cluster respectively.

**SUPPLEMENTARY FIGURE 4.**
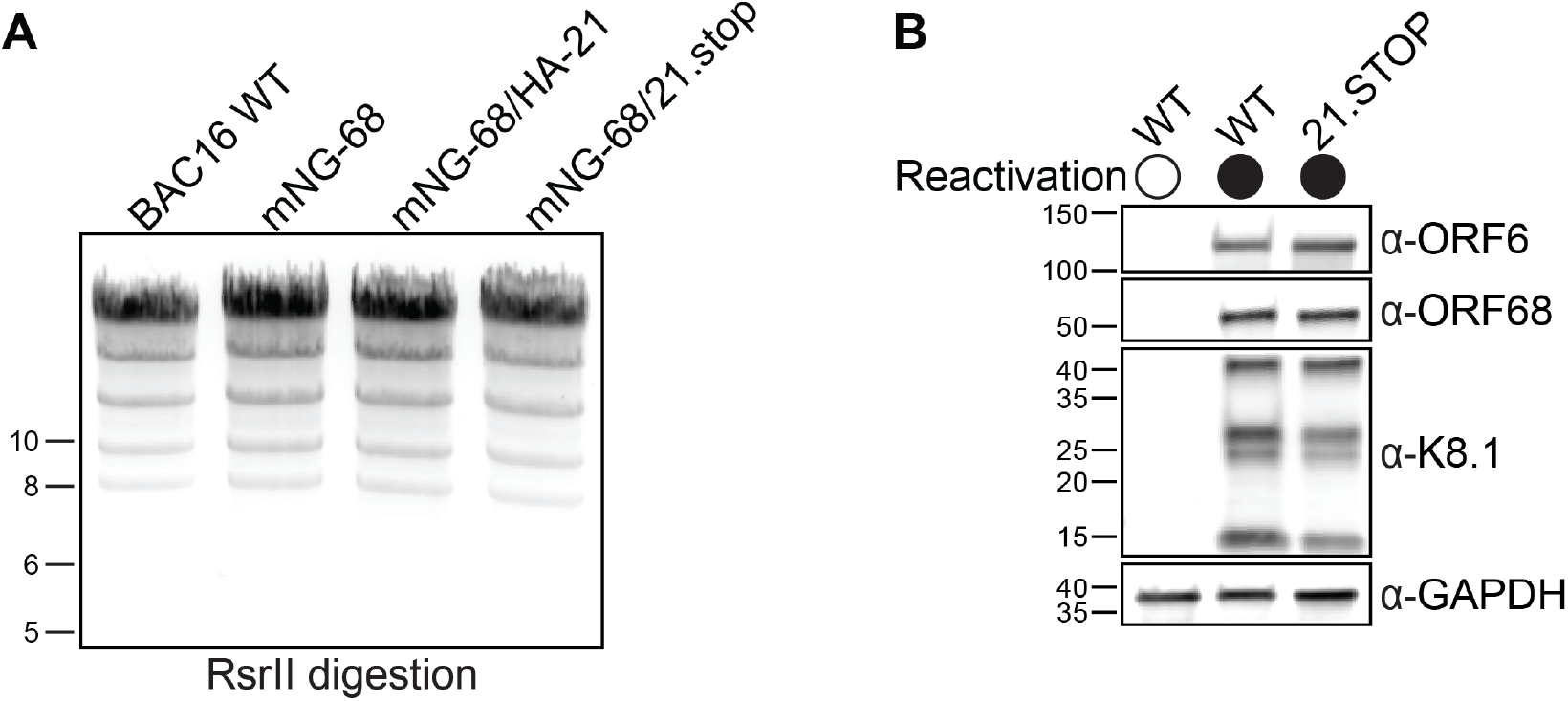
Generation of KSHV cell lines harboring ORF21-HA and ORF21.stop mutations. **A**. Recombinant mNG-ORF68, mNG-ORF68/HA-ORF21, and mNG-ORF68/ORF21.stop BACs were digested with RsrII to assess that no large-scale rearrangements occurred during cloning. **B**. Western blot of whole cell lysate (25 μg) from unreactivated or reactivated WT iSLK cells or iSLK cells harboring the ORF21.stop BAC, representative of 4 biological replicates.

**SUPPLEMENTARY FIGURE S5.**
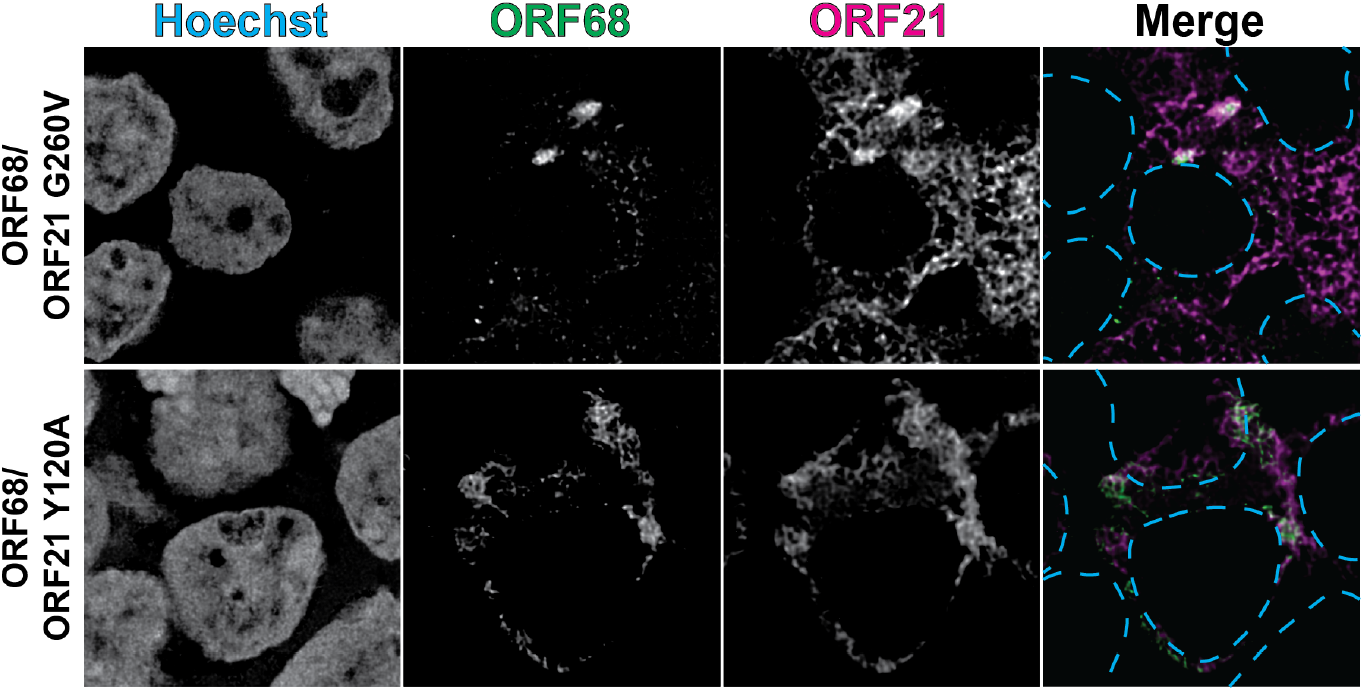
Mutations in the N-terminus and catalytic domain of ORF21 do not alter the localization of ORF68. Z-slices of transfected HEK293T cells expressing HA-ORF68 and full-length ORF21-strep with mutations G260V and Y120A. Stained for Hoechst (cyan dashed lines), HA-tag (green), and Strep-tag (magenta), representative of at least 5 cells from 1 biological replicate. Scale bars are 10 μm.

**SUPPLEMENTARY TABLE 1.** TurboID MS dataset listing all cellular and viral proteins identified across all streptavidin enrichments and replicates.

**SUPPLEMENTARY TABLE 2.**
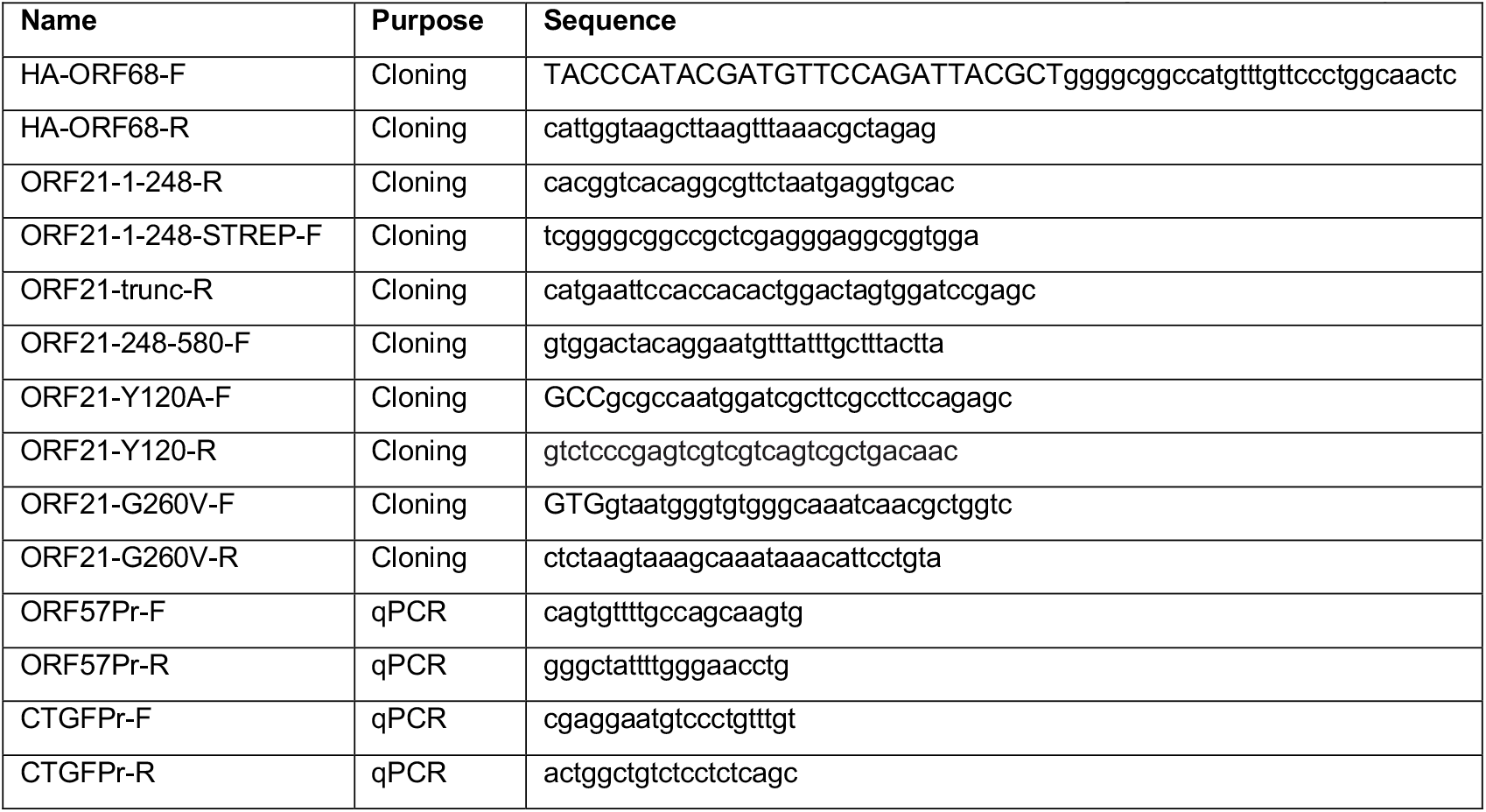
DNA primers used for inverse PCR cloning and qPCR analysis.

**SUPPLEMENTARY TABLE 3.**
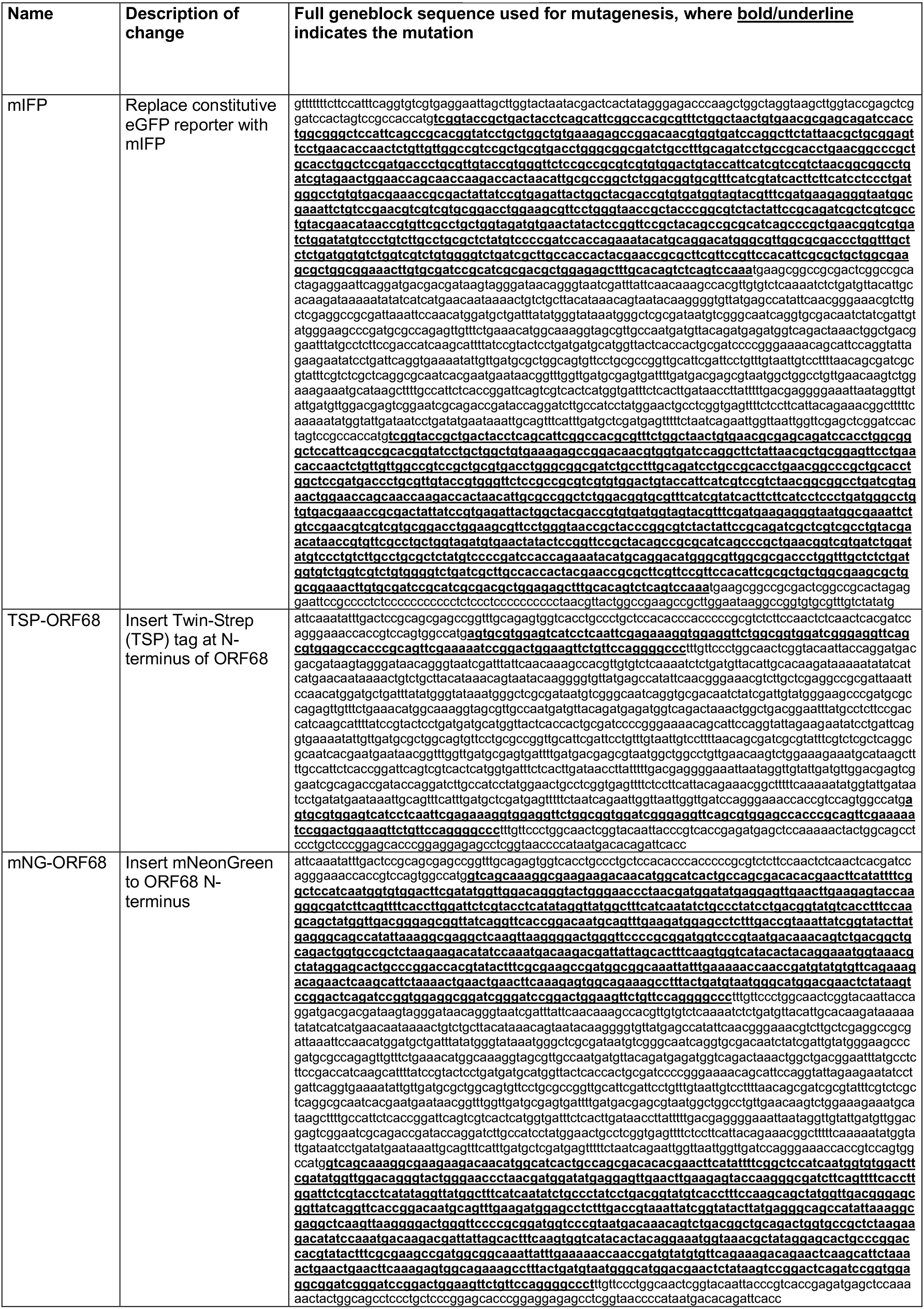
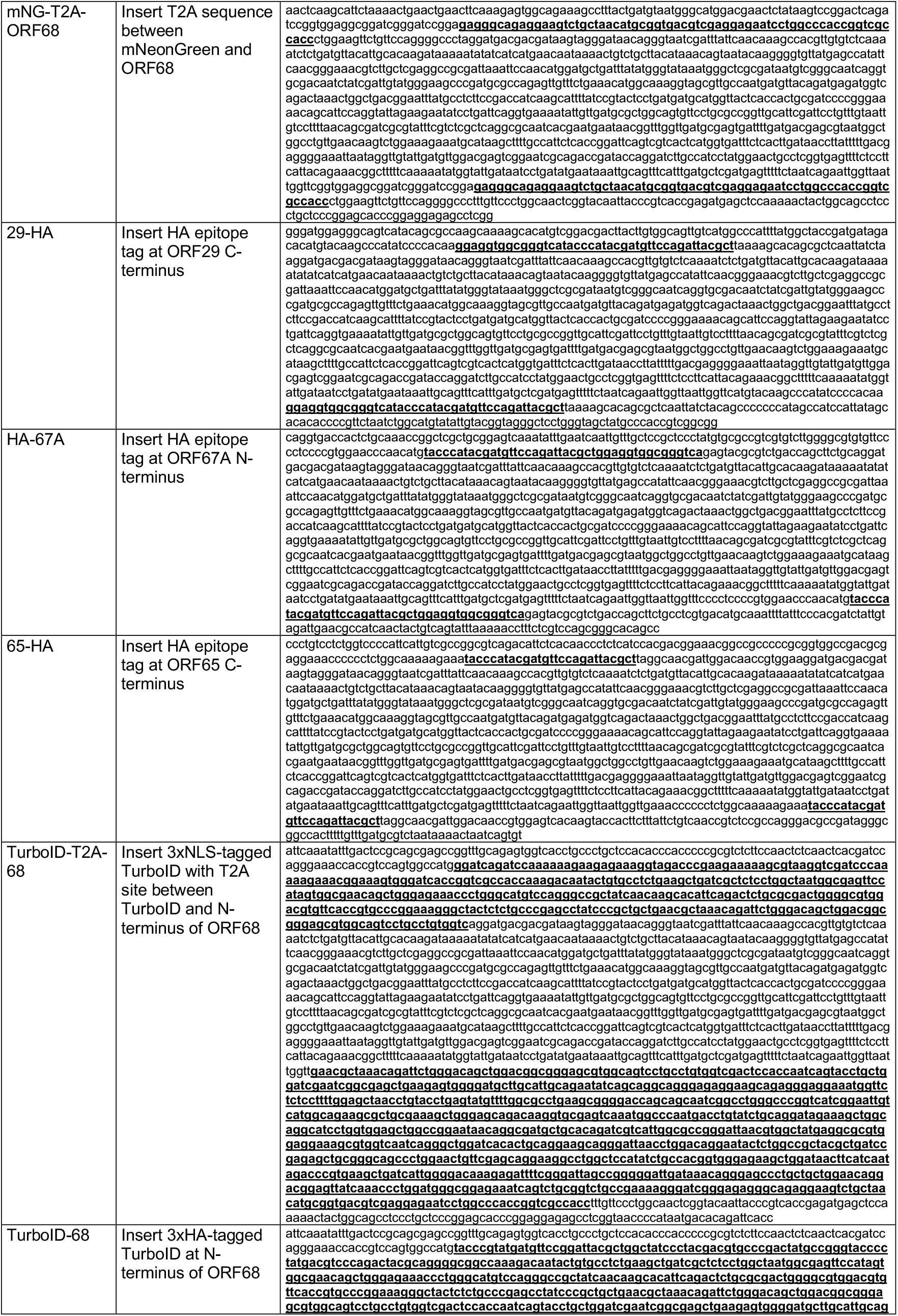
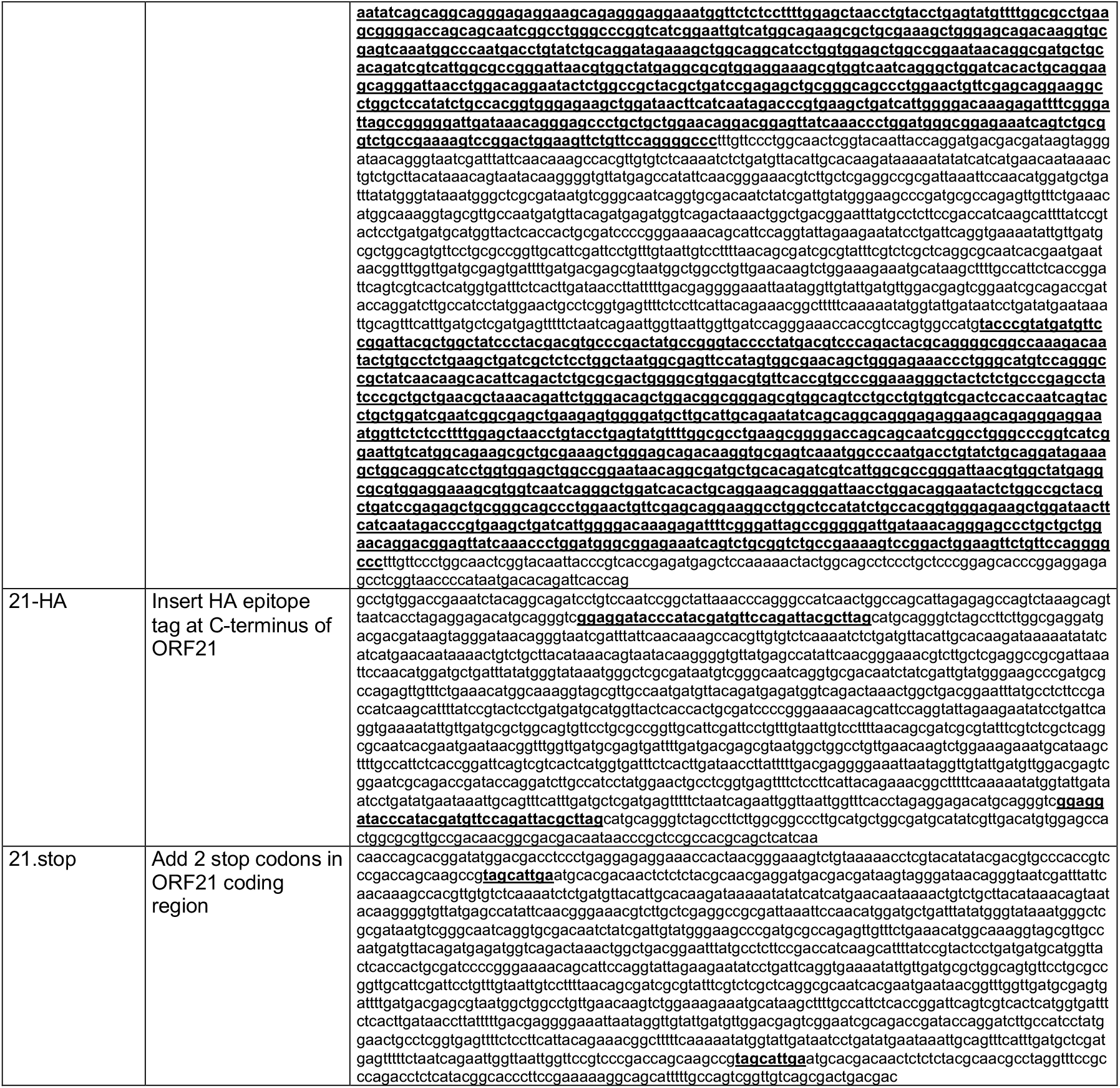
Double-stranded DNA “geneblocks” used for BAC mutagenesis.

**SUPPLEMENTARY TABLE 4.** Primary and secondary antibodies used for western blot and immunofluorescence analysis.

| Antibody | Manufacturer | Part# | Host | Clonality | Use | Dilution |
| --- | --- | --- | --- | --- | --- | --- |
| Vinculin | Abcam | ab91459 | rabbit | polyclonal | Western | 1:1,000 |
| BirA (mutated TurboID) | Agrisera | AS20 4440 | rabbit | polyclonal | Western | 1:5,000 |
| ORF6 | Didychuk et. al 2020<sup>98</sup> | - | rabbit | polyclonal | Western | 1:10,000 |
| ORF26 | US Biological | 524848 | mouse | monoclonal | Western | 1:1,000 |
| ORF68 | Gardner & Glaunsinger 2018<sup>29</sup> | - | rabbit | polyclonal | Western | 1:5,000 |
| K8.1 | Didychuk et. al 2020<sup>98</sup> | - | rabbit | polyclonal | Western | 1:10,000 |
| HA epitope tag | Cell Signaling | 2367S | mouse | monoclonal | Western | 1:1,000 |
| Streptavidin-HRP | Invitrogen | S911 | - | - | Western | 1:500 |
| GAPDH | ThermoFisher | AM4300 | mouse | monoclonal | Western | 1:5,000 |
| Strep tag II | Invitrogen | PIMA537747 | mouse | monoclonal | Western | 1:500 |
| Anti-Strep-tag II | Abcam | ab307676 | rabbit | monoclonal | Western, IF primary | 1:1,000, 1:100 |
| Recombinant Anti-Lamin A + Lamin C | Abcam | ab133256 | rabbit | monoclonal | IF primary | 1:200 |
| HA-Tag (F-7) | Santa Cruz Biotechnology | sc-7392 | mouse | monoclonal | IF primary | 1:500 |
| HHV-8 PF-8 (ORF59) | MilliporeSigma | MABF2752 | mouse | monoclonal | IF primary | 1:200 |
| Nucleolin | Cell Signaling | 14574S | rabbit | monoclonal | IF primary | 1:1000 |
| G3BP1 | Invitrogen | 704023 | rabbit | polyclonal | IF primary | 1:100 |
| PMP70 | Invitrogen | PA1-650 | rabbit | polyclonal | IF primary | 1:200 |
| LAMP1 | Invitrogen | MA5-29385 | rabbit | monoclonal | IF primary | 1:100 |
| Streptavidin, Alexa Fluor™ 568 conjugate | Invitrogen | S11226 | - | - | IF secondary | 1:200 |
| Alexa Fluor® 680 AffiniPure™ Donkey Anti-Mouse IgG (H+L) | Jackson ImmunoResearch | 715-625-150 | donkey | polyclonal | IF secondary | 1:200 |
| Alexa Fluor® 680 AffiniPure Donkey Anti-Rabbit IgG (H+L) | Jackson ImmunoResearch | 711-625-152 | donkey | polyclonal | IF secondary | 1:200 |
| Alexa Fluor® 594 AffiniPure AffiniPure Donkey Anti-Rabbit IgG (H+L) | Jackson ImmunoResearch | 711-585-152 | donkey | polyclonal | IF secondary | 1:200 |
| Alexa Fluor® 594<br>AffiniPure Donkey Anti-Mouse IgG (H+L) | Jackson<br>ImmunoResearch | 715-585-150 | donkey | polyclonal | IF secondary | 1:200 |
| Alexa Fluor® 488<br>AffiniPure Donkey Anti-Mouse IgG (H+L) | Jackson<br>ImmunoResearch | 715-545-150 | donkey | polyclonal | IF secondary | 1:200 |
| Alexa Fluor™ Plus 405<br>Phalloidin | ThermoFisher | A30104 | - | - | IF secondary | 1:400 |

